# A decline in skeletal muscle NOX4 abrogates adaptive homeostasis and exacerbates ageing

**DOI:** 10.64898/2026.03.27.714615

**Authors:** Chrysovalantou E. Xirouchaki, Eamon Coughlan, Esther Garcia-Dominguez, Saveen Giri, Meagan J. McGrath, Florian Wiede, Junichi Sadoshima, William Roman, Maria C. Gomez-Cabrera, Andrew Philp, Marcus Moberg, William Apró, Christina A. Mitchell, Tony Tiganis

## Abstract

A decline in NFE2L2-orchestrated adaptive homeostasis and oxidative distress are thought to be key features of ageing. In contracting skeletal muscle, the ROS-producing enzyme NADPH oxidase 4 (NOX4) is a potent inducer of NFE2L2 adaptive homeostasis. Here we report that skeletal muscle NOX4 levels decline in aged mice and humans, resulting in abrogated NFE2L2 adaptive homeostasis, increased protein oxidative damage and decreased muscle function. We show that deleting NOX4 in skeletal muscle exacerbates the physiological decline associated with ageing, resulting in overt sarcopenia and frailty, characterised by physical inactivity, increased adiposity, systemic inflammation, whole-body insulin resistance and advanced liver disease in aged chow-fed mice. The systems-wide physiological decline in aged skeletal muscle NOX4-deficient mice could be corrected by activating NFE2L2 with sulforaphane and reinstating adaptive homeostatic responses otherwise induced by exercise. Our findings provide important insights into the basis for the decline in NFE2L2-orchestrated adaptive homeostasis with age and identify key mechanisms by which exercise may promote healthy ageing.

## INTRODUCTION

All organisms are constantly exposed to varied internal and environmental stressors, including heat or cold shock, osmotic stress, nutrition, caloric restriction/starvation, hypoxia and exercise training ^1,2^. Adaptations to stressors can increase resilience and allow organisms to manage damaging insults. The ability of organisms to transiently adapt to otherwise harmful stressors is known as adaptive homeostasis ^1,2^. A quintessential example of adaptive homeostasis is the response to oxidants, namely reactive oxygen species (ROS). ROS are highly reactive chemicals generated in response to stressors and as biproducts of life in an aerobic environment ^3,4^. While evolution has harnessed specific ROS, for example H_2_O_2_ for physiological roles such as cellular signaling, stressors resulting in redox imbalance and excess ROS can damage essential macromolecules including proteins, lipids and DNA to promote cell death and inflammation and contribute to disease ^3,4^. The damaging effects of ROS or the inability to cope with ROS-mediated macromolecular damage have long been thought to be important contributors to ageing ^1,2^.

One way by which organisms adapt to ROS is by activating an evolutionary conserved defence system orchestrated by the transcription factor nuclear factor erythroid 2-related factor 2 (NFE2L2) ^4,5^. NFE2L2 activation increases the abundance of hundreds of protective enzymes that limit ROS, ameliorate oxidative damage, eliminate damaged macromolecules and elicit long-lasting protection from subsequent exposures to ROS-producing stressors ^4,5^. NFE2L2 is normally targeted for degradation by the KEAP-1(Kelch-like ECH-associated protein-1)/Cullin-3 E3 ligase complex. However, ROS can oxidise Cys residues (Cys-151, Cys-273, and Cys-288) on KEAP-1 to facilitate NFE2L2’s release and translocation to the nucleus where together with small MAF (musculoaponeurotic fibrosarcoma) subfamily transcription factor it binds to antioxidant response elements (AREs) in the promoter regions of genes ^4,5^. NFE2L2 drives the expression of >200 ARE-containing antioxidant/detoxifying and cytoprotective enzymes ^4,5^. These include enzymes involved in NADPH production necessary for the reduction of oxidised glutathione, including NAD(P)H dehydrogenase (quinone 1) (NQO1) ^6^ as well as the expression of ROS detoxification enzymes, including superoxide dismutase (SOD)-1 and mitochondrial SOD-2 that dismutate superoxide (O2•^−^) into H_2_O_2_, and peroxiredoxins (PRDXs), glutathione peroxidase (GPX)-1 and catalase that eliminate H_2_O_2_ ^4,5^. NFE2L2 also drives the expression of components of the 20S proteosome to facilitate the degradation of damaged proteins ^7,8^. It regulates the cargo recognition protein p62/sequestosome 1 (SQSTM1) that facilitates the autophagic degradation of protein aggregates and organelles ^9^. Additionally NFE2L2 directly and indirectly inhibits inflammation^10^ and supports mitochondrial function, in part by driving the expression of mitochondrial biogenesis genes, including *Pcg1a* (which encodes peroxisome proliferator-activated receptor gamma coactivator 1; PGC-1α) ^11–13^. Growing evidence indicates that ageing is associated with a decline in NFE2L2 activity, oxidative distress and the damage of macromolecules ^14–18^. Indeed, there is a growing appreciation that NFE2L2-orchestrated adaptive homeostasis is compromised during ageing, contributing to the accumulation of damaged proteins, inflammation and the deterioration of cellular function ^1,2,18^.

In mammals, skeletal muscle constitutes 30-50% of body weight and contracting skeletal muscle is a major producer of ROS and a potent inducer of NFE2L2-orchestrated adaptive homeostasis ^19–21^. Such adaptive responses include the promotion of antioxidant defence, mitochondrial biogenesis, glucose uptake/glycogen synthesis and insulin sensitivity ^19–23^. Therefore, adaptative responses to muscle comntraction during exercise serve to bolster respiratory capacity and endurance, afford metabolic flexibility, limit oxidative damage and promote insulin sensitivity to deal with subsequent bouts of intense physical activity. While mitochondria generate the majority of steady state O2•^−^ in muscle cells, NADPH oxidase (NOX)2 and NOX4 are primarily responsible for ROS generation during exercise ^4,20^. NOX2 and NOX4 are differentially localised in skeletal muscle; while NOX2 localises to the plasma membrane of sarcolemma and transverse tubules, NOX4 localises to the sarcoplasmic reticulum, transverse tubules and the inner mitochondrial membrane ^20,24–26^. Although both NOXs can generate O2•^−^, NOX4 is unique in its ability to also directly generate H_2_O_2_ ^27^. The expression of NOX4 and to a lesser extent the catalytic subunit of NOX2 (encoded by *Cybb*) are induced when mice are exercised ^28^. In addition, elegant biosensor studies have shown that NOX2 is activated and generates ROS in contracting skeletal muscle ^29^. The activation of NOX2 has been reported to promote glucose uptake during moderate intensity exercise without overtly altering mitochondrial biogenesis or enhancing insulin sensitivity ^20,29–31^. Using a whole-body loss-of-function model of NOX2 Henriquez-Olguin *et al.* additionally reported that NOX2-deficiency compromises the mitochondrial network remodelling and blunts the improvements in exercise capacity otherwise associated with 6 weeks of high-intensity interval training ^20,32^. By contrast we and others have shown that NFE2L2 adaptive responses in both skeletal and cardiac muscle following acute moderate or high intensity exercise or 5-week exercise training are reliant on NOX4 and the generation of H_2_O_2_ ^20,28,33^. In particular, in skeletal muscle, the induction of NOX4 after exercise is essential for NFE2L2-mediated mitochondrial biogenesis and muscle function/exercise capacity ^28^. It is also plays a critical role in promoting NFE2L2-driven antioxidant defence mechanisms that limit mitochondrial oxidative stress and prevent the oxidative damage of proteins and lipids, which otherwise impairs insulin signaling and promotes insulin resistance ^28^.

We have shown previously that skeletal muscle NOX4 but not NOX2 levels decline in aged mice, or high fat diet fed obese mice, to promote insulin resistance ^28^. In this study we have examined the importance of the decline in NOX4 on NFE2L2 adaptive responses in aged mice and determined the extent to which this might contribute to oxidative distress and the systemic physiological decline associated with ageing.

## RESULTS

### Human skeletal muscle NOX4 and NFE2L2 adaptive homeostasis decline during ageing

To explore the extent to which perturbations in skeletal muscle adaptive homeostasis may contribute to ageing and the decline in physiological integrity in humans, we took advantage of a publicly available RNA sequencing (RNAseq) dataset (GSE159217) assessing the effects of ageing on *vastus lateralis* skeletal muscle gene expression in young (19-25 years old, n=20) versus old (65-71 years old, n=18) male subjects (recruited based on similar physical activity) ^34^. Previous studies using this data set identified oxidative metabolism as the principal pathway downregulated in the muscle of aged individuals ^34^; this was accompanied by the downregulation of mitochondrial biogenesis genes and mitochondrial respiratory proteins for Complexes I and IV ^34^. As NFE2L2 is considered essential for skeletal muscle mitochondrial biogenesis in response to exercise ^11,13,35^ we focussed on pathways regulated by NFE2L2, including the unfolded protein response, ER processing and mitophagy pathways that are induced by NFE2L2 and inflammatory pathway that are repressed by NFE2L2 ^8–10,36^. We found that ageing was associated with an overt repression in NFE2L2 responses, as reflected by the decline in unfolded protein response, ER processing, and mitophagy pathways and the induction of interferon (IFN) α/γ and inflammatory response pathways (**Fig. 1a**; **Fig. S1**). Consistent with this we found that the REACTOME KEAP1/NFE2L2 Pathway gene set was significantly downregulated in the skeletal muscle of aged individuals (NES = -1.643, p < 0.001) (**Fig. 1b**). Furthermore, an analysis of a curated NFE2L2 target gene set focused on antioxidant defence (**Table S1**) revealed that ageing was associated with a decline in skeletal muscle antioxidant defence (NES = -1.615, p = 0.019) (**Fig. 1b**). In particular, genes encoding key antioxidant defence enzymes, including mitochondrial-targeted SOD-2 that eliminates mitochondrial O2•^−^ by converting it to H_2_O_2_ ^37,38^, SOD-1 that similarly eliminates O2•^−^ in the cytosol, GPX-1 and PRDX-1 that eliminate H_2_O_2_ in the cytosol, PRDX-3 that eliminates H_2_O_2_ in mitochondria, and NQO1 that is exclusively regulated by NFE2L2 in muscle ^6^, were decreased in the skeletal muscle of aged subjects (**Fig. 1c**); by contrast the gene encoding catalase (*Cat*) that eliminates H_2_O_2_ at peroxisomes and within the cytosol was not altered (**Fig. 1c**). To ascertain whether reductions in antioxidant gene expression may be associated with decreased antioxidant defence and increased oxidative damage, we took advantage of skeletal muscle *vastus lateralis* biopsies from physically active young (27.6 ± 6.6 years old) and aged (69.9 ± 3.1 years old) men that were matched for body mass index (BMI _young_ = 24.3 ± 0.2; BMI _aged_ = 22.9 ± 0.1) (**Fig. S2a**) and monitored for changes in the abundance of antioxidant enzymes as well as the oxidative damage of proteins (protein carbonylation) by immunoblotting (**Fig. 1d, h**). Consistent with our gene expression analyses we found that SOD-2, PRDX1, PRDX3, GPX1 and NQO1 were downregulated in skeletal muscle lysates of aged men, whereas catalase levels were not altered (**Fig. 1d; Fig. S2b**). Moreover, the decreased abundance of mitochondrial SOD-2 was independently substantiated using mass spectrometry-based proteomic analyses of mitochondria-enriched fractions from hip skeletal muscle biopsies of BMI-matched aged, physically inactive (86.7 ±9.7years old, n=11, 4 men, 7 females; BMI= 26.9 ± 3 kg/m^2^) versus young, physically fit (37.3 ± 10.6 years old, n=8, 5 men, 3 females; BMI= 27± 4.6 kg/m^2^) individuals (**Fig. 1e**; **Fig. S3a**). As a control for mitochondrial content we also monitored for the abundance of the voltage-dependent anion channel 2 (VDAC2) protein that is localised in the outer mitochondrial membrane; VDAC2 levels were similar between young and aged samples (**Fig. 1f**). By contrast mitochondrial complex proteins that are known to decline in the aged were reduced (**Fig. S3b**). In addition, in the whole skeletal muscle lysates from aged men, where we found that the abundance of antioxidant defence enzymes was reduced and that the levels of NOX4 (assessed using a validated antibody^28^), which is required for H_2_O_2_ production and the induction of NFE2L2 adaptive responses in muscle (including mitochondrial biogenesis and antioxidant defence) after exercise, was also reduced (**Fig. 1g; Fig. S2c**). The decreased abundance in skeletal muscle NOX4 and enzymes involved in antioxidant defence was in turn accompanied by increased oxidative damage, as assessed by monitoring for protein carbonylation in the skeletal muscle lysates of aged versus young men (**Fig. 1h**). Taken together these results point towards NOX4 and NFE2L2 adaptive responses, including antioxidant defence, declining in human skeletal muscle during ageing.

**Figure 1.**
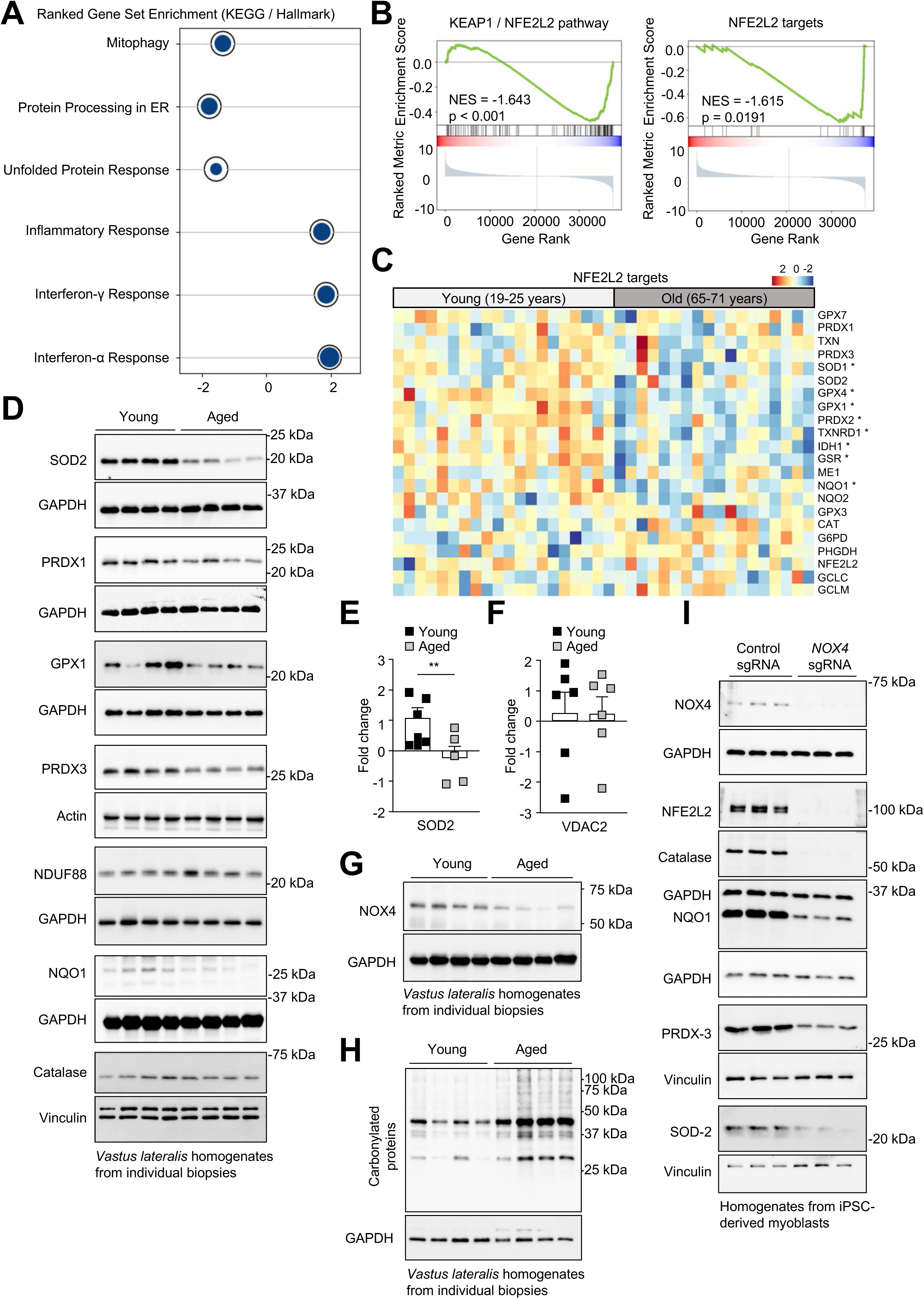
Skeletal muscle NOX4 abundance and NFE2L2 adaptive homeostasis decline during ageing in humans. **a-c)** RNAseq data from GSE159217 was analysed for differential gene expression in human male *vastus lateralis* muscle from aged (65-71 years old, n=18) and exercise-matched young (19-25 years old, n=20) subjects. **a)** Dot plot highlighting significant (p<0.05) negative enrichment of Unfolded Protein Response (Hallmark gene sets) and ER processing and Mitophagy (KEGG Pathways) genes in aged males, and significant (p< 0.05) positive enrichment of Interferon α & γ Response and Inflammatory Response (Hallmark gene sets) genes in aged males. Circle diameters within rings indicate proportions of the reference gene sets present in dataset. **b)** Barcode plots demonstrating negative enrichment of Reactome KEAP1/NFE2L2 Pathway (left) and NFE2L2 curated target set (right) in aged males. **c)** Heatmap of expression levels of the curated set of NFE2L2 targets across the cohort. Genes with significantly downregulated expression (log fold change < 0, p < 0.05) are marked with an asterisk. **d**) Skeletal muscle *vastus lateralis* homogenates from young (27.6 ± 6.6 years old) and aged (69.9 ± 3.1 years old) men that were matched for BMI were analyised by immunoblotting. **e-f**) Mass spectrometry-based proteomic analyses of mitochondria-enriched fractions from hip skeletal muscle biopsies of BMI-matched aged, physically inactive (86.7 ± 9.7 year old) versus young, physically fit (37.3 ± 10.6 years old) individuals to assess **e**) SOD2 and **f**) VDAC protein levels. **g-h**) Skeletal muscle *vastus lateralis* homogenates from BMI matched young versus aged men were assessed by immunoblotting for **g**) NOX4 protein, and **h**) protein carbonylation. **i**) Human iPSCs were differentiated into myoblasts and *NOX4* deleted by CRISPR RNP gene-editing. Cells were processed for immunoblotting. Representative and quantified results are shown (means ± SEM); significance determined using a Student’s t-test (d-i).

Skeletal muscle is a complex tissue consisting of not only multinucleated muscle fibers, but also immune cells, endothelial cells, muscle stem cells and other mononuclear cells important for muscle homeostasis and exercise responses ^39–43^ that change with ageing. Therefore, we sought to determine if the decline in antioxidant defence may be cell autonomous and attributed to the decline in NOX4 in human muscle cells. To this end we took advantage of human induced pluripotent stem cells (iPSCs) and differentiated these into myoblasts as defined by the nuclear expression of the myogenic transcription factor MyoD (**Fig. S4a**) and deleted *NOX4* by CRISPR/Cas9 ribonucleoprotein gene-editing; myoblasts were cultured in the presence of 5% O_2_ to limit oxidative stress (**Fig. 1i**; **Fig. S4b**). The deletion of *NOX4* in myoblasts reduced NOX4 protein by >90% and was accompanied by an overt decline not only in NFE2L2 protein, but also SOD-2, PRDX-3, GPX1 and catalase protein levels **(Fig. 1i**; **Fig. S4c**); mitochondrial content as reflected by the abundance electron transport chain (ETC) complex I protein NDUF88 was not altered (**Fig. S4d)**. The deletion of *NOX4* and the decline in NFE2L2 antioxidant defence was in turn accompanied by the increased oxidative damage of proteins as assessed by monitoring for protein carbonylation (**Fig. S4e**). Therefore, these results are consistent with the apparent decline in NFE2L2 adaptive homeostasis in the skeletal muscle of aged humans being attributed to the decline in NOX4 in muscle cells.

### Skeletal muscle NOX4 and NFE2L2 adaptive homeostasis decline during ageing in mice

To determine whether NFE2L2-orchestrated antioxidant defence may also decline in the skeletal muscle of ageing mice, we monitored for the expression of NFE2L2 target genes by quantitative real time PCR (qPCR) in the *gastrocnemius* muscle of 6-, 12- and 20-month-old male versus female C57BL/6 mice fed a standard chow diet (4.8% fat). Key antioxidant defence genes including those encoding *Nfe2l2*, *Nqo1, Sod1*, *Sod2*, *Prdx1*, *Prdx3* and *Cat* started to decline in both male and female mice by 12-months of age (**Fig. 2a**; **Fig. S5a**); this persisted at 20-months of age for all antioxidant defence genes, with the exception of *Cat* in female mice and *Prdx1* in male mice which increased but was nonetheless lower than that in 6 month old mice (**Fig. 2a**; **Fig. S5a**). The decline in antioxidant defence gene expression was accompanied by a decline in the abundance of NFE2L2, SOD-2, PRDX1, NQO1 proteins as well as a modest albeit not significant decline in catalase levels as assessed by immunoblot analysis in the *gastrocnemius* muscle of male mice (**Fig. 2b**; **Fig. S5b**). Furthermore, we found that skeletal muscle NOX4 protein and/or *Nox4* mRNA also declined by 12-months of age in male and female chow-fed mice and persisted at 20-months of age (**Fig. 2a-b**; **Fig. S5a-b**) Importantly, the decline in NOX4 and NFE2L2 antioxidant defence was in turn accompanied by increasing oxidative damage as reflected by the increasing carbonylation of proteins with age (**Fig. 2c**). Taken together, these results point towards the decline in NOX4 with age abrogating NFE2L2-orchestrated antioxidant defence to promote progressively worsening macromolecular oxidative damage.

**Figure 2.**
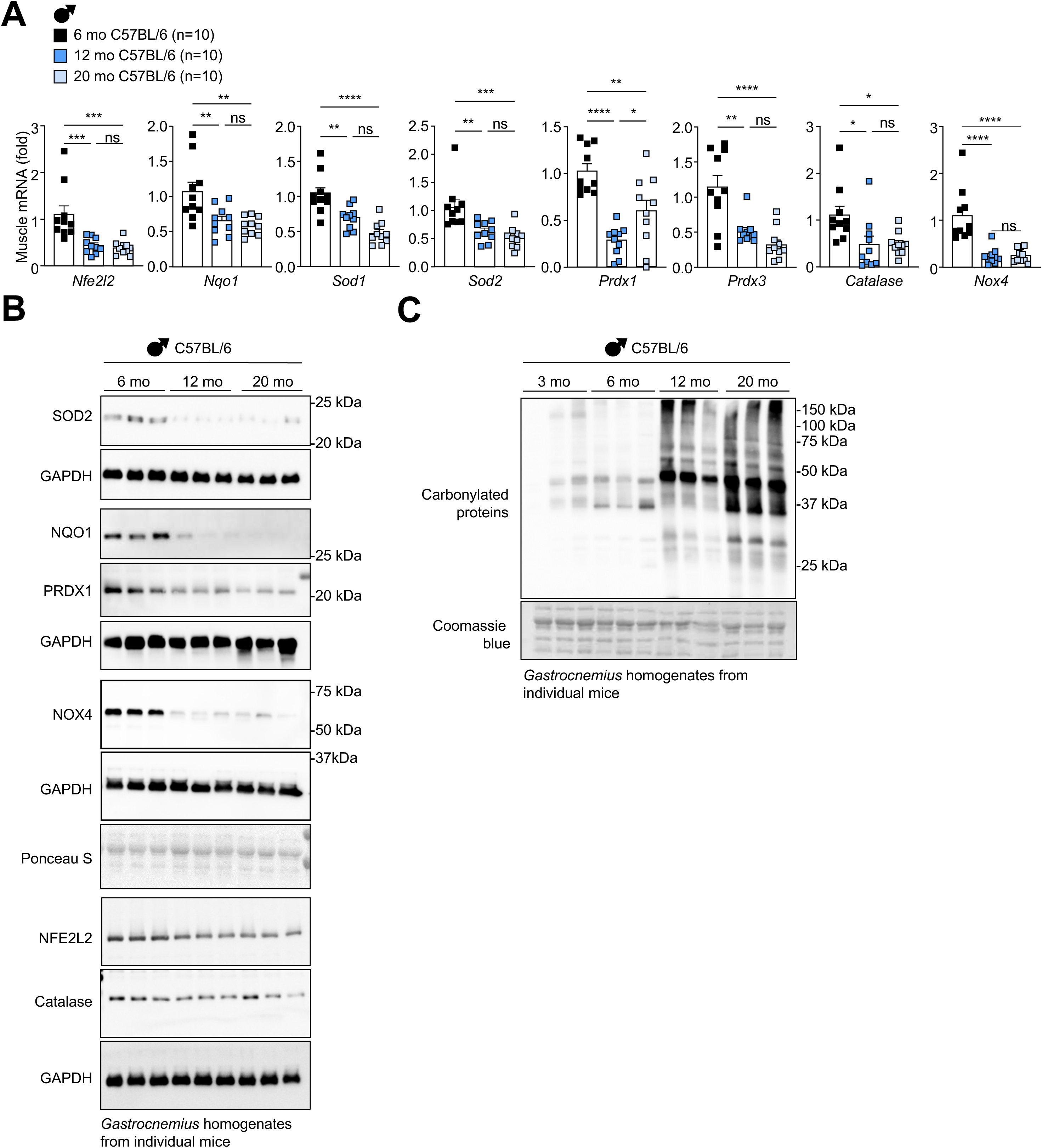
Skeletal muscle NOX4 and anti-oxidant defence decline during ageing in mice. Male C57BL/6 mice were fed a standard chow-diet (4.8% fat) for 3, 6, 12 or 20-months as indicated. *Gastrocnemius* muscle was extracted and analysed by **a**) quantitative real time PCR (qPCR) or **b**) immunblotting. **c**) Alternatively muscles were processed for analysis of protein carbonylation or commassie blue staining. Representative and quantified results are shown (means ± SEM) for the indicated number of mice; significance determined using a one-way ANOVA.

### NOX4-deficiency abrogates NFE2L2 adaptive responses and promotes sarcopenia with age

To explore the extent to which the decline in NOX4 may contribute to the physiological decline associated with ageing, *Nox4*^fl/fl^ control and Mck-Cre;*Nox4*^fl/fl^ muscle-specific NOX4-deficient male and female mice were fed a 4.8% fat chow diet for 6-months or 20-months and effects on ageing assessed; body weights in *Nox4*^fl/fl^ control mice fed a 4.8% fat chow diet for 20 months averaged <40 g. We have reported previously that *Nox4*^fl/fl^ control and Mck-Cre;*Nox4*^fl/fl^ fed a standard chow diet (8.5% fat) for 20 months, or even a high fat diet (23.5% fat; 46% energy from fat) for 20 weeks, so that body weights in each case averaged ∼50 g, do not exhibit any difference in body weights between genotypes ^28^. Consistent with this, body weights and body composition were not altered in 6-month-old *Mck*-Cre;*Nox4*^fl/fl^ versus *Nox4*^fl/fl^ male or female mice fed a 4.8% fat chow diet (**Fig. S6a-b**). By contrast, by 20-months of age, *Nox4*^fl/fl^ male or female mice fed a 4.8% fat chow diet gained less weight than those fed the 8.5% fat diet and male NOX4-deficient mice fed the 4.8% fat chow diet tended to gain more weight whereas the corresponding female mice gained signficantly more weight than *Nox4*^fl/fl^ controls. In both male and female mice this was attributed to increased adiposity (**Fig. 3a-b**; **Fig. S7a-b**), as reflected by body composition analyses (EchoMRI) and white adipose tissue (WAT) weights (inguinal and gonadal) (**Fig. 3b-c**); no differences were evident in lean mass (**Fig. 3b**; **Fig. S7a**) as assessed by EchoMRI (measures smooth, cardiac and skeletal muscles plus all organs). The increased adiposity at 20 months of age was accompanied by decreased voluntary wheel running and energy expenditure (**Fig. 3d; Fig. S7c**); respiratory exchange ratios (RERs; indicative of carbohydrate versus fat oxidation) were modestly reduced in male but not female mice during the night phase (**Fig. 3d; Fig. S7c**). Paradoxically in male mice, NOX4-deficiency was also accompanied by decreased food intake at 6 months and 20 months of age (**Fig. 3d; Fig. S8a**). The decreased food intake coincided with the activation of the integrated stress response (ISR) as early as 6-months of age (**Fig. S8b**). The ISR is a conserved cellular stress response that downregulates protein synthesis and upregulates the expression of specific metabolism-correcting genes in response to stressors, including impairments in mitochondrial function, ER stress and macromolecular oxidative damage ^44^. Central to the ISR is the phosphorylation of eukaryotic translation initiation factor 2 α (eIF2α) on Ser-51 that allows for eIF2α to decrease overall protein synthesis while driving the translation of select genes, such as activating transcription factor 4 (ATF4) ^44^. ATF4 can drive the expression of the transcription factor C/EBP homologous protein (CHOP), DNA damage-inducible protein (GADD34) and secreted factors such as fibroblast growth factor 21 (FGF21) and growth differentiation factor 15 (GDF15) that alter systemic metabolism ^44–47^ to increase energy expenditure and repress feeding respectively ^44,48–51^. We found that eIF2α Ser-51 phosphorylation and CHOP-1, GADD34 and FGF21 protein levels were increased in *Mck*-Cre;*Nox4^fl/fl^ gastrocnemiu*s skeletal muscle (**Fig**. **S8b**) and accompanied by increased FGF21 (**Fig**. **S8c**) and GDF15 in serum (**Fig**. **S8d**). The increased GDF15 in blood in *Mck*-Cre;*Nox4^fl/fl^* mice is consistent with the observed decreased feeding.

**Figure 3.**
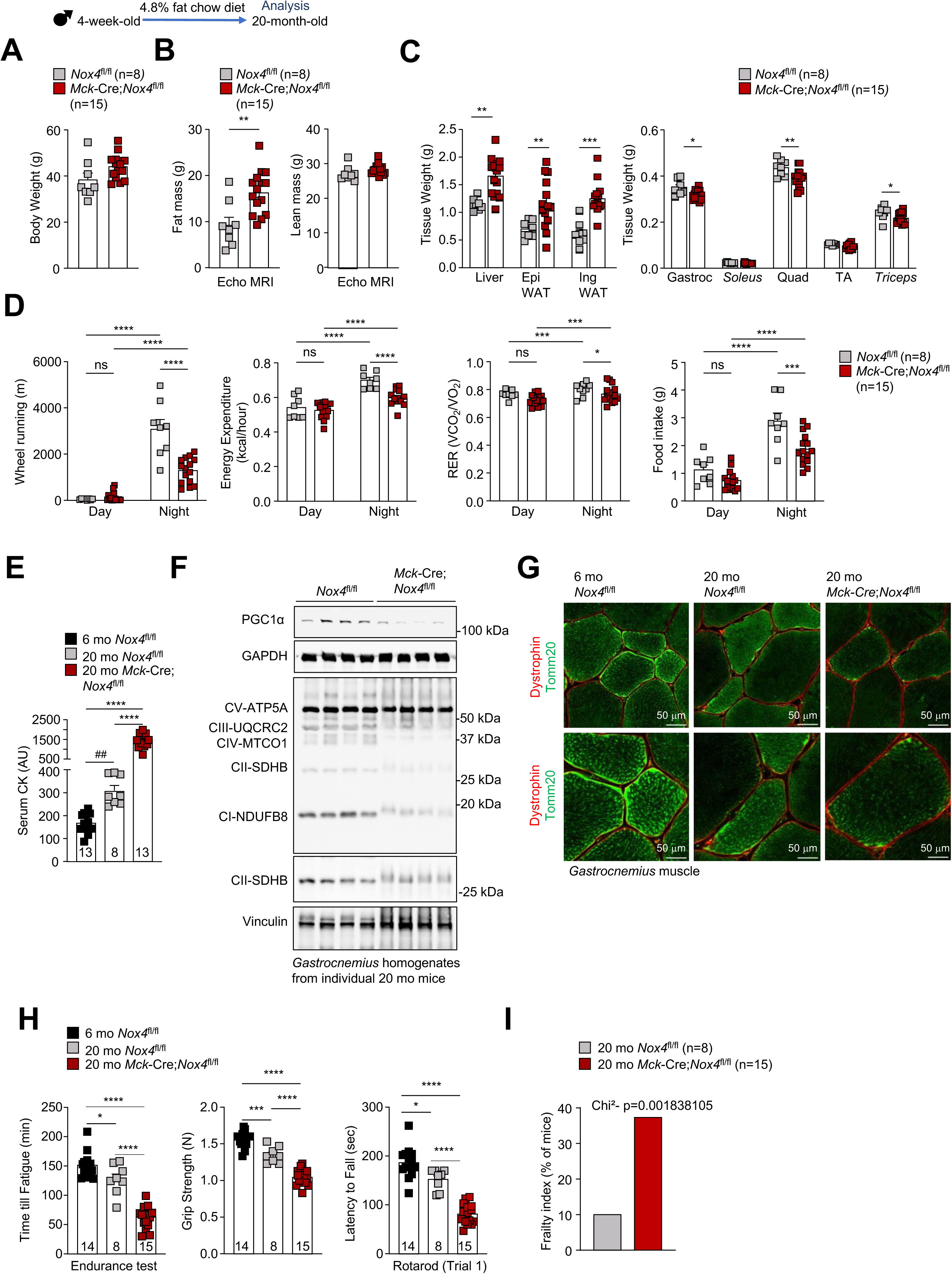
NOX4-deficiency promotes sarcopenia and exacerbates the decline in mmuscle function with ageing. *Nox4*^fl/fl^ control and *Mck*-Cre;*Nox4*^fl/fl^ muscle-specific NOX4-deficient male mice were fed a standard chow diet (4.8% fat) for 20-months. **a**) Body weights, **b**) body composition (Echo-MRI) and **c**) tissue weights [epididymal (Epi) and inguinal (Ing) WAT, *gastrocnemius* (Gastroc), *qaudriceps* (Quad), *tibialis anterior* (TA)] were analysed. **d**) Ambulatory activity (wheel running), energy expenditure, RERs and food intake were analysed in metabolic cages (Promethion). **e-i**) 6 or 20-month old *Nox4*^fl/fl^ and Mck-Cre;*Nox4*^fl/fl^ male mice as indicated were fed a standard chow dieta (4.8% fat) and **E**) serum isolated to assess Creatine Kinase (CK) levels, or *gastrocnemius* muscles dissected and processed for **f**) immunoblotting to monitor PGC1α protein levels and OXPHOS protein complexes, or **g**) immunostaining monitoring for TOMM20 and dystrophin to define mitochondria within individual muscle fibres. **h**) 6 or 20-month old *Nox4*^fl/fl^ and Mck-Cre;*Nox4*^fl/fl^ male mice as indicated were fed a standard chow diet (4.8% fat) and subjected to endurance tests, grip strength measurements, and locomotor activity tests (rotarod). **i**) 20-month-old male *Nox4*^fl/fl^ and *Mck*-Cre;*Nox4*^fl/fl^ mice chow fed mice were subjected to frailty score assessments according to Valencia system. Representative and quantified results are shown (means ± SEM) for the indicated number of mice; significance determined by Student’s t-test (a-c), two-way ANOVA (d), one-way ANOVA (e, h), or a Chi-square test (i); where indicated (#) significance determined by Student’s t-test.

Irrespective, we found that 20-month-old male and especially female NOX4-deficient mice were characterised by decreased skeletal muscle weights; this was evident for different skeletal muscles, including those comprised of oxidative and glycolytic (*gastrocnemius*, *quadriceps*) or predominantly glycolytic [*tibialis anterior* (TA), *triceps*] fibre types (**Fig. 3c; Fig. S7b**); NOX4-deficiency did not decrease *gastrocnemius*, *quadriceps*, TA or *triceps* muscle weights in 6-month-old mice (data not shown). The decreased skeletal muscle weights in aged mice was accompanied by increased circulating levels of creatinine kinase (CK) (**Fig 3e**); CK is abundant in skeletal muscle, but normally low in circulation and its presence can be indicative of muscle damage ^52^. Nonetheless, there were no gross signs of muscular dystrophies as reflected by histology (H&E) and dystrophin staining (marks muscle fibre perimeters) (**Fig. S9a**). Also, no alterations were evident in *gastrocnemius* myofiber size or the abundance of oxidative (expressing type I fibre) and glycolytic (expressing type IIa, type IIb and type IIx fibres) muscle fibres, consistent with previous studies showing that NOX4-deficiency does not alter fibre type abundance in skeletal muscle ^28,53^ (**Fig. S9a-b**). However, NOX4 deficiency was accompanied by an exacerbated decline in mitochondrial biogenesis and content, as reflected by the decline in PGC1α (peroxisome proliferator-activated receptor gamma coactivator 1-alpha) protein that otherwise promotes mitochondrial biogenesis ^23,54^ and the decreased abundance of mitochondrial complex proteins as assessed by immunoblotting (**Fig. 3f**; **Fig. S10a-b**). Consistent with this, histological/immunohistochemical analyses revealed that NOX4 deficiency decreased both subsarcolemmal and intermyofibrilla succinate dehydrogenase (SDH; serves as complex II of the ETC) and Tomm20 (a mitochondrial import receptor protein) staining in *gastrocnemius* muscle (**Fig. 3g**; **Fig. S9c**). Taken together, the decline in muscle mass accompanied by the increased serum CK, along with the decreased mitochondrial content, point towards NOX4 deficiency contributing to the development of sarcopenia, a hallmark of ageing.

Next, we sought to assess the impact of NOX4-deficiency on muscle function (**Fig. 3h**; **Fig. S11**), which along with the decreased muscle mass and mitochondrial function/content is known to worsen with age. We first assessed the impact of NOX4-deficiency on exercise capacity, by subjecting mice to endurance tests till exhaustion. We found that NOX4-deficiency exacerbated the decline at 20 months of age in both male and female mice (**Fig. 3h**; **Fig. S11a**). Similarly, NOX4-deficiency exacerbated the decline in muscle strength, as assessed by measuring forelimb grip strength (**Fig. 3h**; **Fig. S11b**). In addition, NOX4-deficiency exacerbated the decline in motor coordination and balance (as assessed in rotarod tests) in both male and female mice (**Fig. 3h**; **Fig. S11c**). In mice, the following criteria can be used to score frailty: unintentional weight loss, poor endurance (running time), slowness (running speed), muscle weakness (grip strength) and low activity levels (motor coordination) ^55,56^. Together these criteria constitute the Valencia score for frailty and can be used to predict premature ageing ^55,56^. Although we did not observe a decline total body weight in 20-month-old *Mck*-Cre;*Nox4^fl/fl^*mice, this was due to the increased adiposity accompanying the decline in locomotor activity and energy expenditure (**Fig. 3a-d**; **Fig. S7**). Using the change in lean mass between 6-month-old and 20-month-old *Nox4^fl/fl^* versus *Mck*-Cre;*Nox4^fl/fl^* mice rather than total body weight, along with our endurance tests, grip strength tests and rotarod neuromuscular coordination tests in 20-month-old we were able calculate a surrogate frailty score that showed that *Mck*-Cre;*Nox4^fl/fl^* mice exhibited a marked increase in frailty (**Fig. 3i**). Therefore, together our results point towards NOX4 deficiency exacerbating the decline in skeletal muscle function and the development of frailty.

For an unbiased perspective of the impact of NOX4-deficiency on skeletal muscle adaptive responses in 20-month-old aged mice, we subjected *gastrocnemius* muscle from 20-month-old *Mck*-Cre;*Nox4*^fl/fl^ versus *Nox4*^fl/fl^ male mice fed the 4.8% fat chow diet to transcriptomics analysis by bulk RNAseq (**Fig. 4**; **Fig. S12a**). A volcano plot analysis revealed that 1,023 genes were differentially expressed, including 580 genes downregulated and 443 genes upregulated (**Fig. S12a**). GO pathway analysis revealed that genes associated with the response to stressors, including the cellular response to heat, amino acid starvation, starvation, heat and oxidative stress were significantly downregulated (FDR<0.01) and accompanied by the downregulation of the response to unfolded protein, consistent with abrogated NFE2L2 adaptive homeostasis and exacerbated ageing (**Fig. 4a**). In addition, fatty acid oxidation pathways and those involved with the response to insulin otherwise necessary for the provision of fuel during exercise and for replenishing glycogen reserves after exercise were also downregulated (**Fig. 4a**). Similarly, KEGG analysis revealed that numerous pathways normally induced as part of NFE2L2 adaptive homeostasis and downregulated during ageing, were significantly downregulated (FDR<0.01) by NOX4-deficiency in 20-month-old mice; these included genes linked to autophagy, mitophagy and protein processing as well as PPAR and AMPK signalling that are linked to fatty oxidation, exercise-induced mitochondrial biogenesis and glucose uptake (**Fig. 4a**). Consistent with these analyses we found that the KEAP/NFE2L2 pathway (NES = -2.388, p<0.001) and a curated NFE2L2 gene set (**Table S1**) focused antioxidant defence (NES = -1.88, p<0.001) were significantly downregulated (**Fig. 4b-c**). Although we noted variability in select NFE2L2 target genes in aged control mice, this may have been linked to varying physical activity and NOX4/NFE2L2-induced responses, as *Nox4* mRNA levels varied and correlated with the extent of voluntary wheel running in 20-month-old *Nox4*^fl/fl^ mice (R=0.994, p=1.006) (**Fig. S12b**). Irrespective the overt decline in pathways linked to NFE2L2-orchestrated antioxidant defence in the *gastrocnemius* muscle of 20-month-old *Mck*-Cre;*Nox4*^fl/fl^ mice was reaffirmed at the protein level as assessed by immunoblotting for SOD2, NQO1 and catalase (**Fig. 4d**). Furthermore, the exacerbated decline in antioxidant defence accompanying NOX4-deficiency in skeletal muscle was in turn associated with the increased oxidative damage of proteins as reflected by protein carbonylation (**Fig. 4e**). Taken together our findings are consistent with skeletal muscle NOX4-deficiency abrogating NFE2L2 adaptive homeostasis to exacerbate the age-associated oxidative damage of macromolecules, and the decline in muscle mass, mitochondrial content and muscle function otherwise accompanying ageing and the development of sarcopenia.

**Figure 4.**
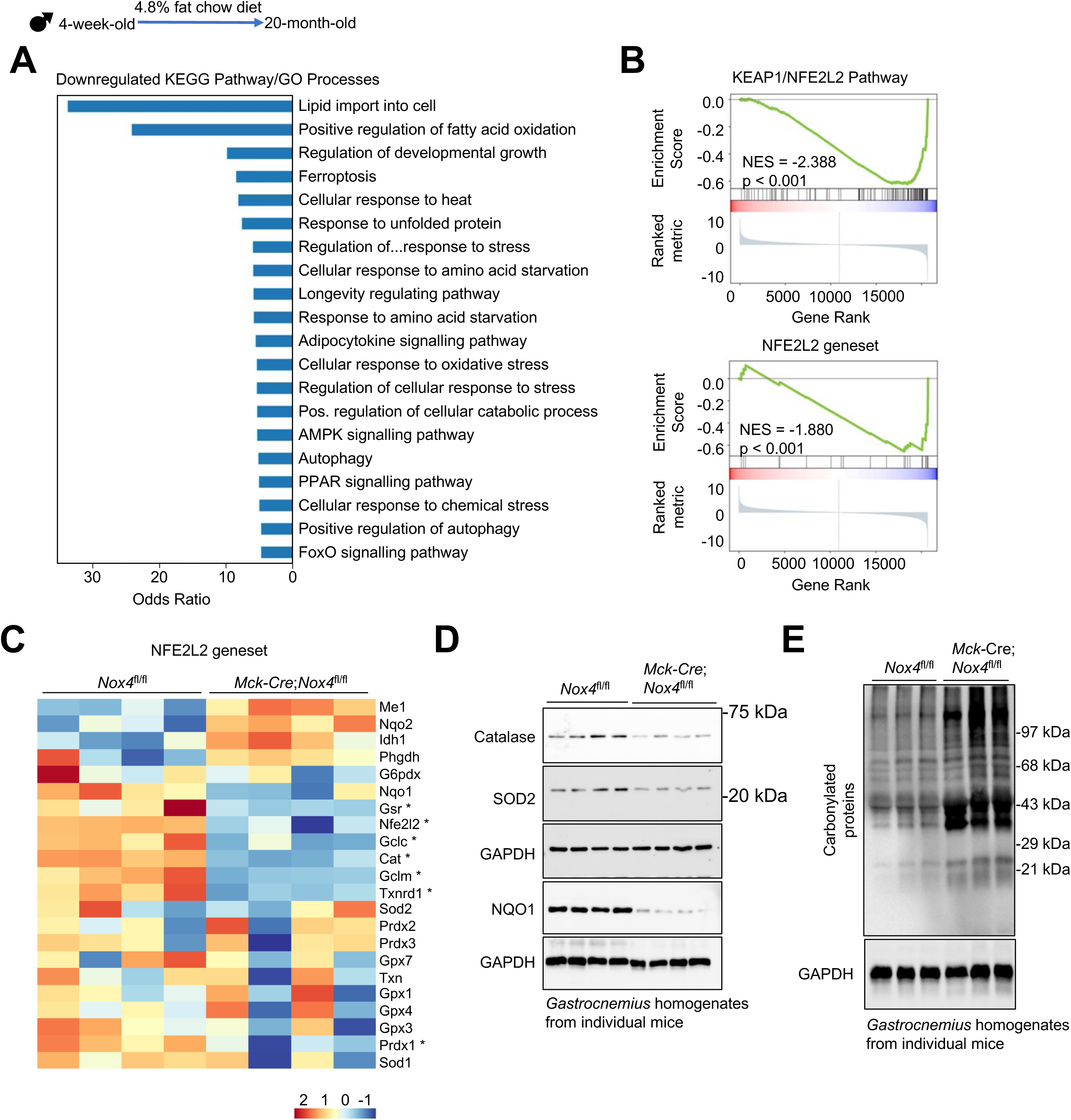
Muscle NOX4-deficiency abrogates NFE2L2-mediated antioxidant defence in aged mice. *Gastrocnemius* from 21-month old *Nox4*^fl/fl^ and *Mck*-Cre;*Nox4*^fl/fl^ male chow (4.8% fat) fed mice (n=4 per genotype) was processed for bulk RNAseq. **a)** Bar plot of Top 20 KEGG pathways/GO Biological Process Terms overrepresented among genes downregulated in *Mck*-Cre;*Nox4*^fl/fl^ mice. Pathways significant with FDR <0.01 were ranked by Odds Ratio. ‘Regulation of…response to stress’ = Regulation of transcription from RNA polymerase II promoter in response to stress. **b)** Negative enrichment of the Reactome KEAP1/NFE2L2 Pathway (top) and NFE2L2 geneset (bottom) in NOX4-deficient animals, demonstrating impairment of ROS-response mimicking that observed in older humans. **c)** Heatmap of NFE2L2 gene set expression. Genes with significantly downregulated expression (log fold change < 0, p < 0.05) are marked with an asterisk. **d**) *Gastrocnemius* muscle from 20-month old *Nox4*^fl/fl^ and *Mck*-Cre;*Nox4*^fl/fl^ male chow (4.8% fat) fed mice processed for immunoblotting to assess SOD2, NQO1 and catalase protein levels and **e**) protein carbonylation. Representative and quantified results are shown.

### NOX4-deficiency promotes systemic inflammation and metabolic disease

A consequence of diminished skeletal muscle NFE2L2 adaptive responses to stressors and ensuing oxidative distress during ageing might be the development of senescence and inflammation ^57–62^. Indeed, previous studies have shown that senescence and inflammatory genes are enriched in the aged and sarcopenic muscles of mice and humans ^57–60^. Consistent with this we found that the decreased muscle mass (**Fig. 3c**) and function (**Fig. 3h**) (sarcopenia) accompanying skeletal muscle NOX4 deficiency in 20-month-old mice was associated with the increased expression of senescence, inflammation and frailty-associated genes (*Gpnmb*, *Spp1*, *Il6*, *Tnf*, *Cxcl1*, *Ccl2*, *Crp*) ^20,58,60,62^ as assessed by qPCR (**Fig. 5a**). An outcome of decrease muscle function/sarcopenia and physical inactivity is increased adiposity, systemic inflammation and metabolic dysfunction. Consistent with this 20-month-old chow-fed *Mck*-Cre;*Nox4*^fl/fl^ mice had significantly increased circulating levels of the proinflammatory cytokine IL-6, with TNF and IFNγ trending higher (**Fig. 5b**) and were overtly hyperglycemic (**Fig. 5c**), hyperinsulinemic (**Fig. 5d**) and insulin-resistant (**Fig. 5e**), as reflected by increased fed and fasted blood glucose and plasma insulin levels and diminished insulin-induced glucose lowering in insulin tolerance tests (**Fig. 5c-e**). Hyperinsulinemic-euglycemic clamps, a gold-standard measure of insulin sensitivity and glucose homeostasis were undertaken in conscious and free-moving 12-month-old male wild type and muscle NOX4-deficient mice prior to any difference in body weight/composition (**Fig. S13a**) and when skeletal muscle NOX4 levels were reduced in wild type mice. These analyses reaffirmed that NOX4-deficient mice were markedly insulin-resistant [as reflected by the reduced glucose infusion rate (GIR) necessary to maintain euglycemia during the insulin-clamp] and this could be largely attributed to reduced glucose clearance [as reflected by the reduced rate of glucose disappearance (RD), a measure of muscle and adipose tissue glucose uptake] (**Fig. S13b-d**). Moreover, as might be expected from the increased adiposity and the development of whole-body insulin resistance, which would be predicted to result in both lipid flux from adipose tissue to the liver and increased liver lipogenesis, 20-month-old chow-fed *Mck*-Cre;*Nox4*^fl/fl^ mice developed metabolic dysfunction-associated fatty liver disease (MASLD) as assessed histologically by monitoring for hepatic lipid accumulation (steatosis) (**Fig. 5f**) and reflected by the increased liver weights in male mce (**Fig. 3c**). This was accompanied by the increased hepatic expression of genes involved in *de novo* lipogenesis, including *Fasn*, *Srebp1* and *Scd1* (**Fig. 5g**). Remarkably, beyond promoting MASLD, muscle NOX4-deficiency promoted the progression to the more advanced metabolic dysfunction-associated steatohepatitis (MASH), which does not otherwise occur in chow-fed mice or indeed obese C57BL/6 mice fed a high fat diet ^63,64^. The progression to MASH in these mice was characterized by presence of hepatic immune infiltrates, as assessed histologically (**Fig. 5f**) and the increased hepatic expression of *Tnf* (**Fig. 5h**); *Ifng* and *Il6* were not altered (**Fig. 5h**). An expected consequence of immune cell recruitment is liver inflammation, damage and the induction of reparative processes that lead to fibrosis. Consistent with this we found that 20-month-old chow-fed *Mck*-Cre;*Nox4*^fl/fl^ mice exhibited overt signs of liver fibrosis, as reflected by Picrosirius red staining (stains collagen; **Fig. 5f**), the increased hepatic expression of fibrosis-related genes, including *Acta2* and *Tgfb*, indicative of hepatic stellate cell activation, the extracellular matrix genes *Fn1* and *Col1a1* (**Fig. 5i**), as well as the significantly increased abundance of hydroxyproline (**Fig. 5j**), a measure of collagen degradation and the severity of fibrosis. Indeed, muscle NOX4-deficiency in 20-month-old chow-fed was accompanied by the development of overt liver damage, as reflected by the significantly increased presence of the liver enzymes AST (aspartate trasnferase) and ALT (alanine transaminase) in serum (**Fig. 5k**). Taken together these results demonstrate that NOX4 deficiency in skeletal muscle can exacerbate the decline in muscle function, promote sarcopenia and the development of systemic metabolic disease accompanied by advanced liver disease, consistent with the onset of frailty.

**Figure 5.**
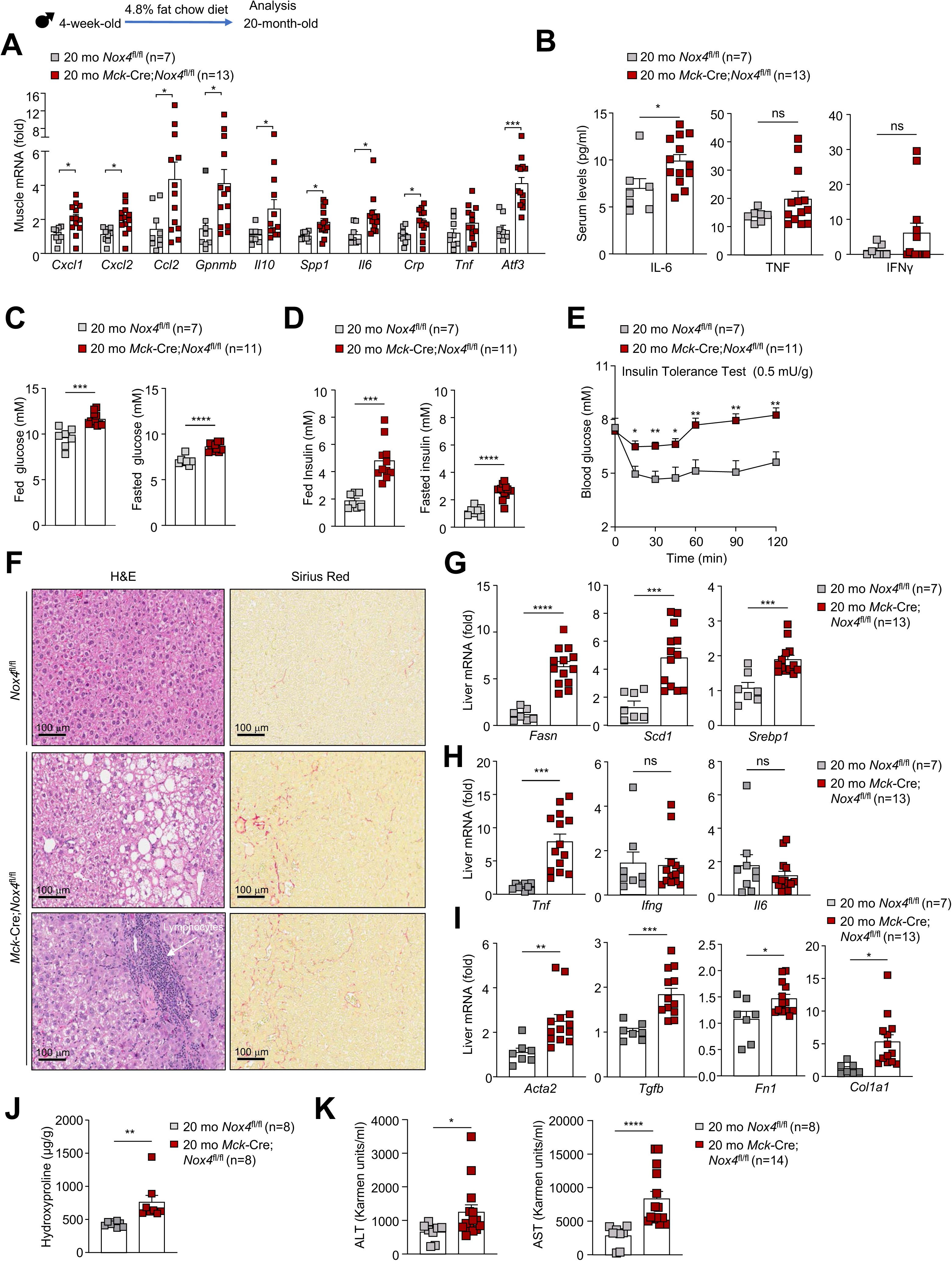
NOX4-deficiency promotes systemic inflammation and metabolic disease in aged mice. *Nox4*^fl/fl^ and *Mck*-Cre;*Nox4*^fl/fl^ male mice were fed a standard chow diet (4.8% fat) for 20-months. **a**) *Gastrocnemius* muscles were extracted and processed for qPCR or blood analysed for **b**) serum IL-6, TNF and IFNγ levels and fed and fasted (6 h) **c**) blood glucose levels, and **d**) plasma insulin levels. **e**) Mice were subjected to insulin tolerance tests (0.5 mg insulin/g body weight). **f**) Livers were processed for histology (H & E and Picrosirius red staining), **g-i**) qPCR to monitor for **g**) lipogenesis, **h**) inflammation and **i**) fibrosis gene expression, or **j**) measurement of liver hydroxyproline levels. **k**) Serum levels of the liver enzymes ALT (alanine transaminase) and AST (aspartate transferase) were assessed. Representative and quantified results are shown (means ± SEM) for the indicated number of mice; significance determined using a Student’s t-test (a-d, h-k) or a two-way ANOVA (e).

### NOX4 is required for the adaptive responses to exercise that mitigate ageing

Having established that skeletal muscle NOX4 and NFE2L2 adaptive responses decline with age and that NOX4-deletion in muscle exacerbates the development of sarcopenia and promotes the onset of frailty, we next sought to determine why NOX4 might decline with age. NOX4 mRNA and protein levels in skeletal muscle started to decline in C57BL/6 mice by 12 months of age (**Fig. 2a-b**). At 12 months of age the decline in NOX4 in skeletal muscle was accompanied by diminished ambulatory activity and voluntary wheel running when compared to 6-month-old mice (**Fig. S14**). We have reported previously that NOX4 levels are induced by exercise in skeletal muscle and that this is required for the induction of NFE2L2-mediated antioxidant defence and mitochondrial biogenesis. Therefore, we asked if the decline in NOX4 may be attributed to the decreased physical activity or otherwise represent an immutable aspect of ageing. To this end, we assessed NOX4 levels and the induction of NFE2L2 antioxidant defence genes in the *gastrocnemius* muscle of 6-month versus 12-month-old sedentary or 12-month-old exercise-trained mice; trained mice were subjected to 5 consecutive weeks (3 days/week) of treadmill running progressively increasing intensity from 50% to 80% of their maximum pre-training exercise capacity in each successive week (**Fig. 6a**). We found that NOX4 mRNA and protein levels in *gastrocnemius* muscle in 12-month-old trained mice were reinstated to those in 6-month-old mice (**Fig. 6b-c**). Moreover, we found that exercise training also increased the expression of *Nfe2l2*, *Sod2* and *Nqo1* levels as assessed by qPCR so that they approximated or exceeded those in 6-month-old sedentary mice (**Fig. 6b**). In addition, exercise training ameliorated the increased oxidative damage of proteins otherwise associated with ageing in *gastrocnemius* muscle (**Fig. 6d**). Therefore, the decline in NOX4 and NFE2L2 antioxidant defence in skeletal muscle can be attributed to the diminished physical activity that occurs with ageing. Importantly, exercise training also enhanced muscle function and motor coordination, as reflected in treadmill endurance tests, grip strength measurements and rotarod performance tests, however, muscle NOX4 deficiency completely abrogated such exercise-induced responses (**Fig. 6e; Fig. S15a; Fig. S16a**). NOX4-deficiency also prevented the induction of NFE2L2 target genes (*Nfe2l2*, *Sod2* and *Nqo1*) (**Fig. 6f**) otherwise associated with exercise training without impacting on body weight or body composition (**Fig. S15b-c; Fig. S16b-c**). Therefore, these results ascribe the ageing-associated decline in NOX4 to the decline in physical activity and establish that muscle NOX4 is essential for NFE2L2 adaptive responses to exercise. Moreover, our findings are consistent with the ageing-associated decline in NOX4 being causal in the accompanying decline in physiological integrity.

**Figure 6.**
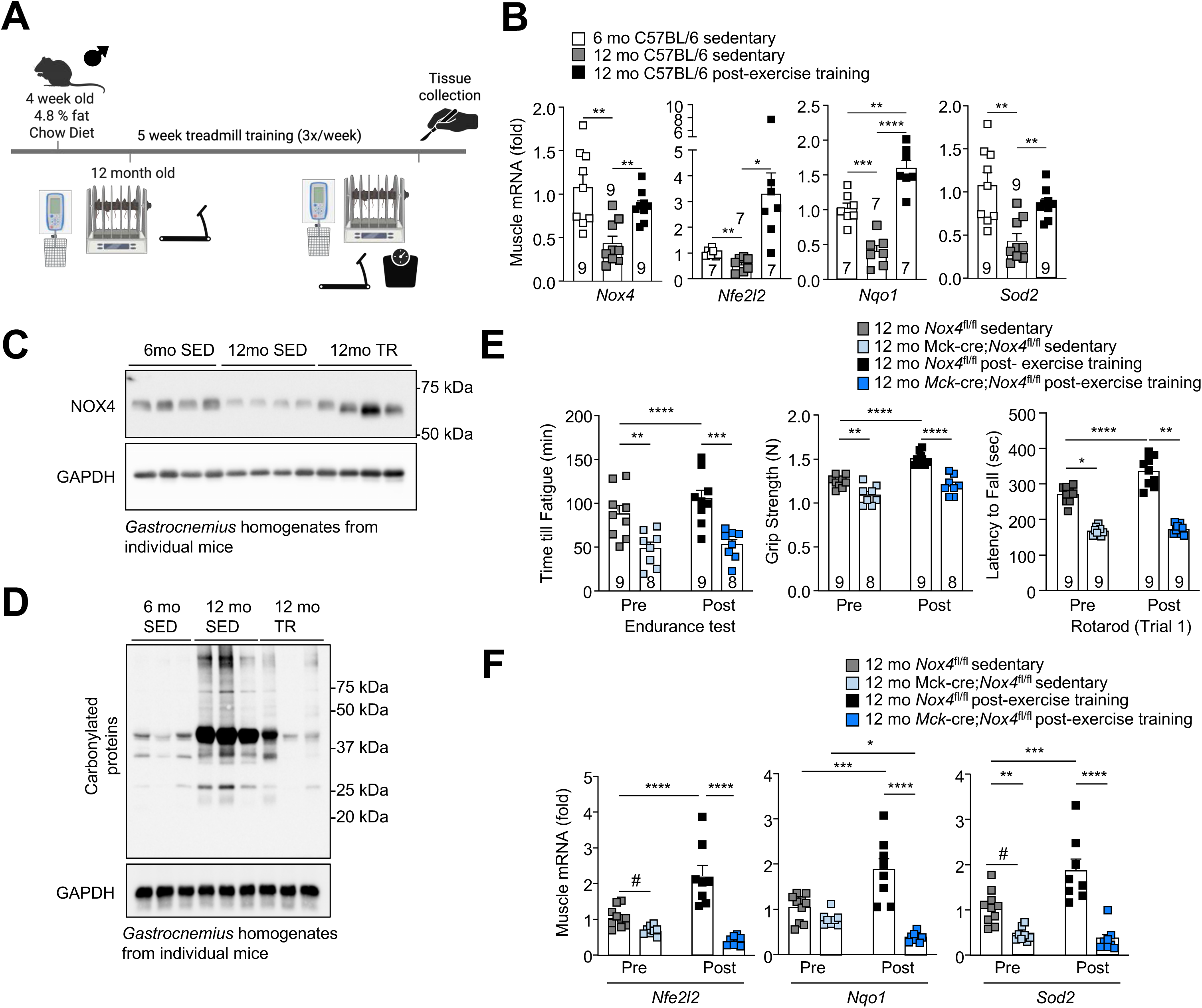
NOX4 is required for the adaptive responses to exercise. C57BL/6 mice were fed a standard chow diet (4.8% fat) for 6 or 12 months and were either exercise trained (TR) for 5 weeks (with progressively increasing running speeds each week) or left sedentary (SED). **a**) Schematic representation of the exercise training and subsequent analyses. **b-d)** *Gastrocnemius* muscles were extracted and processed for **b**) qPCR or **c-d**) immunoblotting to monitor for **c**) NOX4 or **d**) protein carbonylation. **e-f**) *Nox4*^fl/fl^ and *Mck*-Cre;*Nox4*^fl/fl^ male mice were fed a standard chow diet (4.8% fat) for 12 months and subjected to basal and post exercise training (5 weeks) and measurements of endurance, grip strength and locomotor activity (rotarod; Trial 1). **f**) Alternatively, mice were either left sedentary or trained for 5 weeks and gastrocnemius muscle extracted for qPCR. Representative and quantified results are shown (means ± SEM) for the indicated number of mice; significance determined using a one-way ANOVA (b), two-way ANOVA (e-f) or where indicated (#) by Student’s t-test.

### NFE2L2 activation tempers ageing in muscle NOX4-deficient mice

Physical activity and capacity progressively decline with age and the implementation of regular exercise regimes in older people with increasing frailty and/or in people with diverse ethnic and socioeconomic backgrounds remains a significant challenge. An alternative approach may be to take advantage of NFE2L2 agonists to mimic the NFE2L2 adaptive homeostatic responses that are induced by exercise and decline with age. The isothiocyanate sulforaphane is naturally occurring NFE2L2 agonist that is abundant in cruciferous vegetables and has been reported to protect skeletal muscle and other tissues from oxidative damage ^65,66^. Accordingly, we asked if sulforaphane could reverse the sarcopenia and frailty evident in our aged muscle-specific NOX4-deficient mice. 21-month-old 4.8% fat chow-fed *Mck*-Cre;*Nox4*^fl/fl^ mice were administered vehicle or sulforaphane (2 mg/kg, 3x/week, i.p.) for 4 weeks and effects on body weight, muscle function, metabolic health and NFE2L2 adaptive responses assessed (**Fig. 7**). Remarkably, sulforaphane reversed the increased body weight (**Fig. 7a**) and adiposity (**Fig. 7b-c**), otherwise evident in 22-month-old vehicle-treated *Mck*-Cre;*Nox4*^fl/fl^ mice and conversely increased muscle mass to that in 22-month-old *Nox4*^fl/fl^ control mice (**Fig. 7c**). Indeed, the improved muscle mass was accompanied by the correction of serum CK otherwise at pathological levels in 22-month-old vehicle-treated *Mck*-Cre;*Nox4*^fl/fl^ mice, to those in 22-month-old *Nox4*^fl/fl^ control mice (**Fig. 7d**). This was accompanied by the reinstatement of NFE2L2 orchestrated antioxidant defence, as reflected by expression of NFE2L2 target genes (*Nfe2l2*, *Nqo1*, *Cat*, *Sod2*) (**Fig. 7e**) and the corresponding NFE2L2, SOD-2 and catalase protein (**Fig. 7f**) and the attenuated oxidative damage of proteins (**Fig. 7g**) in *gastrocnemius* muscle. More importantly, sulforaphane corrected the otherwise decreased *gastrocnemius* muscle mitochondrial content (as assessed by histologically monitoring for Tomm20 staining) (**Fig. 7h**) and decreased exercise capacity and motor coordination (measured by rotarod tests) (**Fig. 7i**) and the overt metabolic dysfunction associated with muscle NOX4-deficiency in aged mice (**Fig. 7j-o**). In particular, sulforaphane corrected the hyperglycemia (**Fig. 7j**), the increased hepatic (*Tnf*) and systemic inflammation (as reflected by the elevated serum TNF and IL-6 levels) (**Fig. 7k-l**) and the increased liver weights (**Fig. 7c**) and steatosis, as assessed histologically (**Fig. 7m**) and reflected by the repressed expression of the lipid synthesis genes *Fasn* and *Scd1* and the trending repression of *Srebp1* (**Fig. 7n**). Sulforaphane treatment also corrected the ongoing liver damage as reflected by the attenuated serum levels of AST and ALT (**Fig. 7o**), but had no impact on the extent of fibrosis (Picrosirius red staining) (**Fig. 7m**) that is known to be progressively irreversible. Taken together these results suggest that NFE2L2 agonists can reinstate NOX4/NFE2L2 adaptive homeostatic responses in skeletal muscle to temper the physiological decline otherwise associated with physical inactivity in the aged.

**Figure 7.**
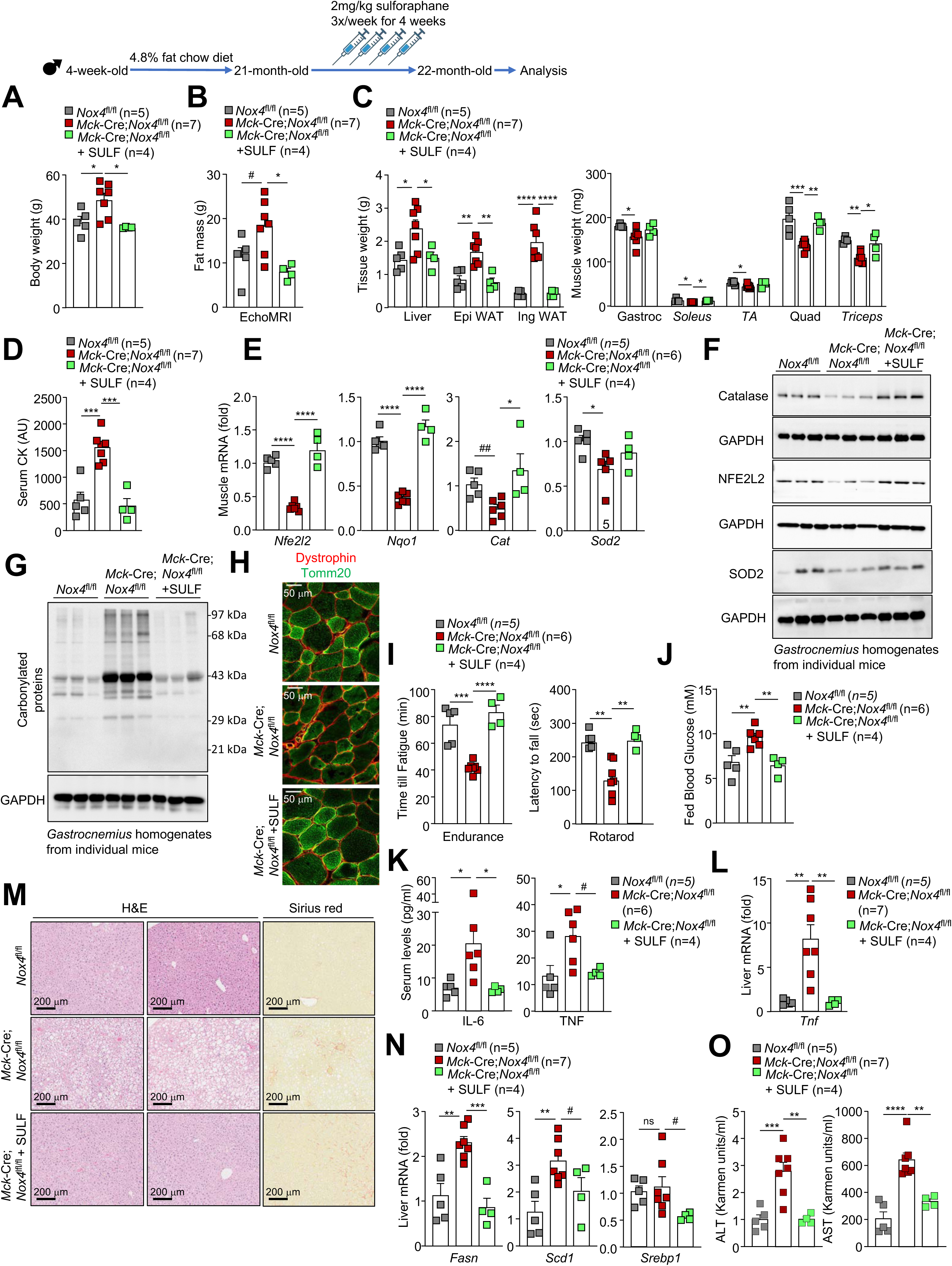
Sulforaphane treatment corrects the physiological decline and metabolic dysfunction in aged muscle NOX4-deficient mice. *Nox4*^fl/fl^ control and *Mck*-Cre;*Nox4*^fl/fl^ muscle-specific NOX4-deficient mice were fed a standard chow diet (4.8% fat) for 21-months and treated with vehicle (DMSO) or sulforaphane as indicated for 4 weeks. **a**) Body weight, **b**) fat mass (EchoMRI) and **c**) tissue weights were assessed. **d**) Serum CK levels were assessed and *gastrocnemius* muscles extracted and processed for **e**) qPCR or **f-g**) immunoblotting monitoring for **f**) NFE2L2, SOD2 and catalase levels or **g**) protein carbonylation, or **h**) immunostaining monitorng for mitochondria (Tomm20) within muscle fibres (dystrophin). **i**) Mice were subjected to tests for endurance or locomotor activity. **j**) Fed blood glucose levels. **k**) Serum IL-6 and TNF levels or **l**) hepatic *Tnf* mRNA were assessed to minitor for systemic and hepatic inflammation respectively. **m**) Livers were processed for histology to monitor for steatosis (H&E) or fibrosis (Picrosirius red). **n**) Livers were processed for qPCR to assess the expression of lipogenesis genes. **O**) Serum ALT and AST levels were measured to monitor for ongoing liver damage. Representative and quantified results are shown (means ± SEM) for the indicated number of mice; significance was determined using one-way ANOVAs (a-o) or where indicated (#) by Student’s t-test.

## DISCUSSION

The evidence supporting the importance of physical activity in promoting resilience and healthy ageing is irrefutable ^22^. Indeed, lifelong physical inactivity in mammals can accelerate the loss in bone density, skeletal muscle mass, muscle strength and power, otherwise associated with ageing, whereas physical activity builds resilience and fitness ^67,68^. The results of this study are consistent with the benefits of physical activity/exercise in promoting healthy ageing being at least in part attributed to NOX4-dependent skeletal muscle ROS generation for the induction of NFE2L2-orchestrated adaptive homeostatic responses that among other things mitigate macromolecular oxidative damage and perturbations in mitochondrial function. Our studies indicate that a diminution in physical activity with age results in a decline in skeletal muscle NOX4 levels, abrogating NFE2L2 adaptive homeostasis thereby resulting in compromised muscle function, increased oxidative macromolecular damage and the progressive development of sarcopenia and frailty. Importantly, our studies suggest that the physiological decline associated with physical inactivity and abrogated NOX4/NFE2L2 adaptive homeostasis can be corrected by the administration of NFE2L2 agonists.

Previous studies have shown that both cardiac and skeletal muscle NOX4 levels are induced in response to exercise training and/or acute exercise in mice ^28,33^. Skeletal muscle *Nox4* mRNA is also induced after acute high intensity interval training in humans ^28^. These studies have shown that the induction of NOX4 and the resultant ROS generation activate NFE2L2 to promote antioxidant defence and mitochondrial biogenesis to 1) enhance cardiac or muscle function and exercise capacity, 2) mitigate the oxidative damage of macromolecules, including proteins and lipids, and 3) temper the resultant mitochondrial oxidative stress and the decline in insulin sensitivity that otherwise occurs with ageing ^28,33^. Yet other studies suggest that NOX4 in the vasculature may also facilitate exercise capacity by promoting glucose and fatty acid oxidation ^69^ or by increasing capillary density in response to exercise ^70,71^ and that NOX4 may also protect the vasculature from ischemic or inflammatory stress ^70^. Recently we have also shown that heightened hepatocyte NOX4 levels in MASLD similarly induce NFE2L2 adaptive responses in hepatocytes to limit oxidative damage, metabolic dysfunction and the progression from simple steatosis to the more advanced MASH with fibrosis ^64^. Herein we have shown that skeletal muscle NOX4, NFE2L2 and downstream targets that function to mediate antioxidant defence decline in abundance in aged mice and humans and that this is accompanied by the oxidative damage of proteins. In mice we show that skeletal muscle NOX4 mRNA and protein levels start to decline by ∼12-months of age when locomotor activity and voluntary wheel running decline, and this coincides with diminished NFE2L2 antioxidant defence and conversely a marked increase in oxidative protein damage that progressively worsens with age. We show that the decline in NOX4 is causally linked with the decline in NFE2L2-orchestrated adaptive homeostasis in several ways. First, the deletion of NOX4 in human iPSC-derived myoblasts is accompanied by the ablation of NFE2L2 protein, a decline in the abundance of key antioxidant defence enzymes, including SOD-2, PRDX-3, NQO1 and catalase, and a marked increase in protein carbonylation. Second, we show that whereas skeletal muscle NOX4, NFE2L2 and NFE2L2 downstream targets decline by 12 months of age, both can be rescued by exercise training in wild type mice, but not in muscle NOX4-deficient mice. Third we show that adaptive homeostatic responses induced by exercise, including enhanced running capacity, muscle strength (grip strength) and motor coordination (rotarod test), are completely abrogated by muscle NOX4-deficiency. Finally, we show that whereas muscle NOX4-deficiency exacerbates the decline in NFE2L2 orchestrated antioxidant defence, the oxidative damage of proteins and the development of sarcopenia/frailty, these can be largely, if not completely corrected with the administration of the NFE2L2 agonist sulforaphane. Taken together, our studies ascribe the increasing oxidative damage of macromolecules, worsening muscle function and the overall decline in physiological integrity that accompany physical inactivity during ageing to the decline in NOX4-instigated and NFE2L2-orchestrated adaptive homeostasis.

Although in this study we have focused on NFE2L2-orchestrated antioxidant defence, the decline in physical activity and NOX4 undoubtedly affect numerous other NFE2L2-orchestrated adaptive responses during ageing. Our previous studies have shown that skeletal NOX4-deficiency reduces mitochondrial biogenesis and content and compromises exercise capacity and endurance ^28^. Consistent with this, others have shown that global NOX4 deficiency attenuates the induction of PGC1α and mitochondrial biogenesis induced by voluntary wheel running ^72^. Similarly, in this study, we show that muscle NOX4-deficiency in aged mice exacerbates the reduction in PGC1α and mitochondrial content associated with physical inactivity during ageing, consistent with a decline in mitochondrial biogenesis. The induction of mitochondrial biogenesis is a fundamental hormetic response to exercise and is considered essential for the promotion of respiratory capacity and endurance ^23,54^. NFE2L2 is required for the induction of mitochondrial biogenesis after exercise and for optimal exercise performance ^11,54,65,66^. Indeed, previous studies have shown that the global deletion of NFE2L2 not only results in the oxidative damage of macromolecules in the muscles of aged mice ^35^ but also in the decline of mitochondrial content and the onset of sarcopenia and frailty ^13^. Several studies have shown that skeletal muscle mitochondrial capacity and/or mitochondrial biogenesis and ETC complex genes decline in aged individuals, however this can be attenuated at least in part by increasing or maintaining physical activity during ageing ^34,61,73–76^. Consistent with this, we found that even in aged NOX4-deficient mice with overt sarcopenia and frailty, the NFE2L2 agonist sulforaphane was able to bypass NOX4-deficiency to increase muscle mass, mitochondrial content and physical activity to mitigate sarcopenia and frailty.

Beyond the promotion of oxidative distress and reduced mitochondrial content, NOX4-deficiency in aged mice additionally promoted local inflammation, as reflected by the increased expression of both senescence and inflammatory genes in skeletal muscle. This may be an outcome of mitochondrial dysfunction and oxidative distress or otherwise the reduced PGC1α that normally inhibits the expression of proinflammatory genes such as TNF and IL-6 ^10,36^. In addition, the decline in NFE2L2-may also contribute, since NFE2L2 can directly repress NFκB expression as well as the expression of proinflammatory cytokines such as IL-6 and IL-1β in a cell context-dependent manner ^10,36^. Alternatively, it is possible that the decline in NOX4/NFE2L2-orchestrated adaptive responses and resultant oxidative distress in multinucleated muscle cells may affect resident mononuclear cells, including muscle stem cells, mesenchymal progenitors, T cells and macrophages that are important for muscle homeostasis, repair and exercise responses ^39–41,77,78^ and whose activation and relative abundance in skeletal muscle changes in aged mice to increase the expression of proinflammatory and senescence-related markers ^58,60^. Irrespective, we propose that the local skeletal muscle inflammation likely contributes to the systemic inflammation and the associated development of both muscle wasting and insulin resistance evident in our NOX4-deficient mice ^78,79^.

Previous studies have reported that skeletal muscle ageing is associated with markers of oxidative damage in muscle tissue and in isolated muscle fibres ^80–82^. Yet other studies suggest that skeletal muscle ageing in mice is associated with a fundamental remodelling of the redox landscape that involves the increased oxidation of proteins linked with muscular dystrophies ^83^. A fundamental question that arises from our studies is how can the decline in NOX4-derived ROS contribute to oxidative distress and the increased oxidation of specific redox networks? One possibility is that this may be attributed to the compartmentalisation of ROS generation during ageing. Studies have shown that mitochondrial, but not cytosolic H_2_O_2_ may be increased in skeletal muscle fibres and in isolated mitochondria from old mice ^15,84,85^. By contrast NOXs, in particular NOX2 and NOX4, rather than mitochondria, are responsible for the beneficial exercise-induced H_2_O_2_ generation required for adaptive responses ^25,28,29,86^. Consistent with this we have shown previously that the deletion of NOX4 in myoblasts and the resultant reduction in H_2_O_2_ is accompanied by increased mitochondrial ROS generation and the oxidative damage of proteins that can be corrected by activating or stabilising NFE2L2, or by administering mitochondrial-targeted antioxidants ^28^. Moreover, we have shown previously that skeletal muscle NOX4 deficiency in 6-month-old mice was accompanied by marked reductions in both SOD-2 and PRDX3 in mitochondria and increased macromolecular damage^28^. Our current studies indicate that SOD2 is similarly reduced in the skeletal muscles of aged mice and that this is reinstated by exercise training or exacerbated by the deletion of NOX4. Numerous studies have shown that mitochondrial oxidative stress and the ensuing oxidative damage contribute to the development of insulin resistance ^35,37,87–90^ and that *Sod2* heterozygosity in mice is sufficient to promote mitochondrial oxidative stress and insulin resistance whereas SOD-2 overexpression attenuates insulin resistance ^37^. Accordingly, we propose that the decline in NOX4-instigated and NFE2L2-orchestrated antioxidant defence with age, which is exacerbated by the deletion of NOX4, result in the decline in SOD2 and PRDX-3 levels to promote oxidative distress and damage and the ensuing insulin resistance. The decreased mitochondrial content, physical activity and the resultant increased adiposity associated with the decline in NOX4 that are also exacerbated by NOX4 deficiency undoubtedly also contribute to the accompanying systemic disease. However, the mechanisms that may contribute to the unexpected progression from simple steatosis to MASH and fibrosis in our aged muscle NOX4-deficient mice remain unclear. Although this may be ascribed to the accompanying systemic inflammation, other factors, including for example changes in myokine/exerkine secretion ^91^ may also contribute, but this requires further investigation.

Our studies suggest that the promotion of NOX4 expression in skeletal muscle in the aged may help mitigate sarcopenia and frailty. However, it is important to recognise that the overexpression or induction of NOX4 in the context of disease can somewhat paradoxically also contribute to oxidative distress and exacerbate pathology. For example, although NOX4 drives adaptive responses in the liver to attenuate MASLD progression ^64^ its upregulation in the context of MASH has been reported to exacerbate disease progression ^92^. Similarly, NOX4 is induced in the failing heart and NOX4 deletion attenuates whereas transgenic NOX4 overexpression exacerbates cardiac hypertrophy, fibrosis and cardiomyocyte apoptosis under conditions of pressure overload induced hypertrophy ^93,94^. As such, in this study we sought to activate NFE2L2 downstream of NOX4 to reinstate the adaptive homeostatic responses that otherwise decline with ageing. Our studies show that the naturally occurring NFE2L2 agonist sulforaphane can bolster antioxidant defence and completely correct the muscle wasting, increased adiposity and diminished muscle function, as well as temper the metabolic perturbations including MASLD, in aged muscle NOX4-deficient mice. Consistent with this, previous studies have shown that the administration of sulforaphane to 21-22-month-old C57BL/6 mice over 3 months enhances muscle function, increases antioxidant defence and improves glucose homeostasis ^95^ whereas the administration of sulforaphane as a concentrated broccoli sprout extract once daily for 12 weeks to obese patients with dysregulated type 2 diabetes (HbA1c 57.1 ± 6.6 mmol/mol) on metformin, lowers HbA1c levels without adverse effects ^96^. Thus, activating NFE2L2 with such agonists may be beneficial for tempering sarcopenia and frailty, especially in aged and physically inactive individuals where exercise may not readily feasible or practical.

Taken together the results of this study reaffirm the fundamental importance of NOX4 in the beneficial adaptive responses induced by physical activity/exercise and demonstrate that physical inactivity and declining NOX4 levels with age are instrumental in abrogating NFE2L2-orchestrated adaptive homeostatic responses that otherwise mitigate the development of sarcopenia and frailty. Our studies provide insight into fundamental mechanisms contributing to the physiological decline with ageing and define an approach for reinstating otherwise abrogated adaptive homeostatic responses that promote healthy ageing.

## ACKNOWLEDGMENTS

We thank YaoYao Jia, Joe Roberts, and Crystal Stivala or technical support and Joel Eliades (Monash Metabolic Phenotyping Platform) for assistance with metabolic phenotyping. This work was supported by the National Health and Medical Research Council of Australia (to T.T.).

## AUTHOR CONTRIBUTIONS

T.T. conceived and conceptualized the study, designed experiments, wrote the manuscript and interpreted the data with intellectual input from all authors. C.E.X. conceptualized and designed experiments, conducted and analysed experiments the majority of experiments and contributed to the reviewing and editing of the manuscript; E.C. undertook bioinformatic analyses; S.G. and M.J.M were responsible for the muscle immunohistochemistry and histomorphometry; E.G.-D. contribute to cell culture experiments and analyses human mitochondrial enriched fraction analyses. Other authors performed and/or analysed experiments. All, authors contributed to the reviewing/editing of the manuscript.

## MATERIALS AND METHODS

### Materials

Mouse α-tubulin (Cat #T5168, RRID:AB_477579), α-vinculin (Cat #V9131, RRID:AB_4776298) and rabbit α-NFE2L2 (Cat #AV38745, RRID:AB_1854419) used for immunoblotting were from Sigma-Aldrich (St Louis, MO); mouse α-NQO1 (Cat #NB200-209, RRID:AB_10002706) from Novus Biologicals (Littleton, CO) and sheep α-NQO1 from R&D Systems (Minneapolis, MN); α-catalase (Cat 1877, RRID:AB_302649), α-PRDX3 (Cat # AB222807) and α-NOX4 (Cat # ab133303, RRID:AB_11155321) were from Abcam (San Francisco, CA); mouse α-GAPDH (Cat #AM4300, RRID:AB_2536381) from Thermo Fisher Scientific (Waltham, MA); rabbit α-catalase (Cat #ab1877, RRID: AB_302649), α-NFE2L2 (Cat #sc365949, RRID:AB_10917561) and α-NOX4 (Cat #sc-30141, RRID:AB_2151703) from Santa Cruz Biotechnology (Santa Cruz, CA) and α-SOD2 (Cat #13141, RRID:AB_2636921) from Cell Signaling Technology (Beverly, MA). Goat α-rabbit IgG (AQ132P) and goat α-mouse IgG (AQ502P) secondary antibodies for immunoblotting were purchased from Merck Millipore (Burlington, Massachusetts, USA) and donkey α-sheep IgG (HAF016) was purchased from BIotechne (Minneapolis, USA). Mouse α-dystrophin (ab7164), rabbit α-dystrophin (ab15277) and rabbit α-Tomm20 (ab78547) used for tissue immunofluorescence microscopy were from Abcam (San Francisco, CA); mouse α-myosin heavy chain I (BA-f8), IIa (SC-71), IIb (BF-F3) and IIx (6H1) were from Developmental Studies Hybridoma Bank (All fiber-typing antibodies were developed by S. Schiaffino (Universita degli Studi di Padova, Italy) or C. Lucas (University of Sydney, Australia) and obtained from the Developmental Studies Hybridoma Bank, created by the NICHD of the NIH and maintained at The University of Iowa, Department of Biology, Iowa City, IA 52242). The Mouse GDF-15 ELISA Kit (ab216947), Hydroxyproline Assay Kit (ab222941), Mouse ALT ELISA Kit (ab282882), Mouse AST ELISA Kit (ab263882) and OxyBlot™ Protein Oxidation Detection Kit (S7150) were from Abcam, (Cambridge, UK) and Rat/Mouse FGF-21 Quantikine ELISA Kit (EZRMFGF21-26K) and Citrate Synthase Assay Kit (MAK116-1KT) from Merck Millipore (Burlington, Massachusetts, USA). Legendplex assays [Mouse IL-6 Flex Set (B4) (558301), Mouse TNF Flex Set (C8) (558299), Mouse IFNγ Flex Set (558296)] were purchased from BD Bioscience (New Jersey, USA).

### Mice

Age-matched male and female murine models were utilized in all experimental conditions. Animals were housed under a controlled 12 h light-dark cycle in a high-barrier, temperature-regulated facility (Monash Animal Research Laboratory-Monash ARL) with unrestricted access to food and water. Mice were fed a standard chow diet (20% protein, 4.8% fat, 4.8% crude fiber; Specialty Feeds). *Mck*-Cre;*Nox4*^fl/fl^, have been described in previous studies ^28,93^. All mice were on a C57BL/6 background. C57BL/6J mice were purchased from Monash Animal Research Platform or Walter and Eliza Hall Institute of Medical Research (WEHI). Experiments were approved by the Monash University School of Biomedical Sciences Animal Ethics Committee (14368, 22138) and were carried out in accordance with the NHMRC Australian Code of Practice for the Care and Use of Animals. All animals were aged to 3-, 6-, 12-, 20- and 22-months for the different aging timeline studies.

### Human participants and biopsies

For immunoblotting *vastus lateralis* muscle from overnight fasted young (27.6 ± 6.6 years old, n=10) and aged (69.9 ± 3.1 years old, n=10) men that were matched for body mass index (BMI _young_ = 24.3 ± 0.2; BMI _aged_ = 22.9 ± 0.1) and had active lifestyles have been described previously ^97,98^ and conformed to the West Midlands – Black Country Research Ethics Committee (#17/WM/0068) and the Swedish Ethical Review Authority (#2017/1139-31/44; #2017/2107-31/2) and adhered to the principles set by the Declaration of Helsinki.

For the proteomic analyses of human mitochondrial fractions, twelve patients (six young adults and six older adults) undergoing trauma or orthopaedic surgery were included in this study. Their medical and functional status was evaluated using the modified five-item Frailty Index ^99^. Exclusion criteria encompassed conditions such as myopathy, hemiplegia or hemiparesis, rheumatoid arthritis or other autoimmune connective tissue disorders, inability to provide consent, hospital admission within the past month, or major surgery in the preceding three months. Each participant provided written informed consent following a comprehensive explanation of the study’s purpose and potential risks. This research adhered to the principles outlined in the Declaration of Helsinki and received ethical approval from the Ethical Committee for Drug Research of the Department of Health Arnau de Vilanova – Liria, Spain (Licence reference: CEIm 28/2019). Skeletal muscle samples were collected during surgery from a healthy muscle region, ensuring the absence of contusion or haematoma. A small portion of the muscle was excised using a scalpel, following the natural alignment of the muscle fibres while avoiding electrocautery. The samples were immediately frozen in liquid nitrogen for subsequent mitochondrial isolation and proteomic analyses.

### Rodent exercise studies

Mice were acclimatised to treadmill running for 3 days prior to the initiation of the experiments. Mice were placed on a multi-lane treadmill (Columbus Instruments, Columbus, OH) for 10 min, mice and run for 5 min at 10 m/min and 1 min at 15 m/min at 0% slope. All animals were randomized prior to the initiation of exercise tests.

#### Exercise stress test

To assess peak oxygen consumption (VO_2 peak_) male mice were placed in an enclosed single lane treadmill connected to Oxymax O_2_ and CO_2_ sensors (Columbus Instruments, Columbus, OH). On the experimental day, prior to the initiation of the exercise stress test, mice were acclimated in the chamber at rest for 20 min (basal O_2_ consumption and CO_2_ generation assessed for the last 5 min). The mice were then subjected to running at 10 m/min at 0% incline and the running speed (U) increased by 4 m/min every 3 min until the mice reached fatigue; fatigue was determined as the time point where the mice could not be prompted to continue running for at least 5 sec. O_2_ consumption (VO_2_max) was assessed at the maximal exercise speed (Umax). Differences between the exercised and basal states were calculated and gas exchange data used to determine energy expenditure, heat and the respiratory exchange ratio (RER = VCO_2_/VO_2_).

#### Endurance test

Mice were exercised on a multi-lane treadmill (Columbus Instruments International) at 0% slope. The endurance was performed with a warmup period of 10 min at 10 m/min, followed by running at 60% of the maximal exercise speed (Umax) until fatigue was reached. The time till exhaustion was considered the representative measure of endurance capacity.

#### Exercise training

Mice were allocated into sedentary and training groups. The training group underwent exercise training for 5 weeks, 3 days per week, while the sedentary group remained sedentary on the treadmill for the same period of time. Mice were subjected to a gradual overload protocol, in which running speed was increased so that by the end of training weeks 1, 2, 3, 4 and 5 mice were running for up to 45 min at 50%, 60%, 70%, 80% and 90% of their maximum pre-training speed respectively. The maximal exercise speed and VO_2_ max were assessed using exercise stress tests.

### Muscle performance tests

#### Grip strength measurement

Limb grip strength was assessed using a grip strength meter (Bioseb, EB Instruments, France). Mice were acclimatised for 3 days by placing all four limbs on the grip strength meter grid, 3 times per day. The acclimatisation was separated by 3 days from the experimental period. On the experimental day, mice were placed over the meter so that all paws grasped the grid while mice were handled by their tails and tails were pulled horizontally until mice released their hold of the grid. Three independent measurements were undertaken and limb strength measured in Newtons.

#### Rotarod test

Mice were trained once per day for 4 days on a rotating rod with a lane width of 5 cm (Ugo Basile Rota-Rod 476000, France) spinning at 4 rpm for 5 min. After training, mice were subjected to an incremental protocol, where the speed was increased over 480s from 4 to 60 rpm. All animals were subjected to four independent trials separated by 1 h and the latency to fall (length of time that the mice remained on the rod) recorded and analysed.

### Metabolic and blood measurements

Mouse body weights were monitored weekly, and body composition was evaluated using EchoMRI (Echo Medical Systems, Houston, TX), as previously described ^28^. Insulin and glucose tolerance tests were carried out following established protocols ^28^. Blood samples were collected from conscious mice in both fed state (satiated, 11 pm add refs) and following a 6-hour fasting period via submandibular bleeding. Plasma insulin concentrations were quantified using the Mouse Insulin ELISA (ALPCO, Salem, 80-INSMS- E01), while corresponding blood glucose levels were measured using an Accu-Chek glucometer.

Energy balance parameters, including food intake, voluntary wheel running activity, and energy expenditure, were continuously recorded over a 72-hour period following a 24-hour acclimatization phase. These assessments were performed using the Promethion Metabolic Screening System (Sable Systems International, NV), equipped with indirect open-circuit calorimetry, running wheels, and automated sensors for quantifying food consumption and locomotor activity.

Hyperinsulinemic-euglycemic clamp studies were conducted as previously described^28^. Mice were anesthetized using 2% (v/v) isoflurane (250 mL/min O₂), and surgical catheterization of the left common carotid artery and right jugular vein was performed to enable arterial blood sampling and intravenous infusions. On the day of the experiment, food was withdrawn, and following a 3.5-hour fasting period, a primed (2 min, 0.5 μCi/min) continuous infusion (0.05 μCi/min) of [3-³H]-glucose was administered to assess basal glucose turnover. After 5 h of fasting, a continuous insulin infusion (4 mU/kg/min) was initiated to achieve a hyperinsulinemic state, while blood glucose concentrations were maintained at euglycemic levels through a dynamic infusion of a 50% (w/v) glucose solution. Arterial blood samples were collected at baseline and at steady-state time points (80, 90, 100, 110, and 120 minutes). At the end of the experiment, mice were euthanized, and tissues were rapidly harvested, flash-frozen in liquid nitrogen, and stored for downstream analyses of gene expression and glucose uptake.

### Frailty assessment: Valencia Score

The “Valencia Score” was used to assess frailty in 20 month old *Mck*-Cre;*Nox4*^fl/fl^ mice^55^. This approach was adapted from the frailty criteria originally developed for humans by Linda Fried and colleagues ^100^. The score includes the measurement of five key components: unintentional weight loss (for which we modified the original protocol and used lean mass loss as the criterion), reduced activity levels (motor coordination), weakness (grip strength), poor endurance (total running time in the incremental treadmill test), and slowness (maximum running speed in the incremental treadmill test). For running time, running speed, and grip strength parameters, we classified as frail the 20% of mice with the lowest performance. This allowed us to determine the percentage of mice considered frail for each parameter. For the body weight criterion, mice were classified as frail if they lost more than 5% of their lean mass compared to the previous measurement. In the case of motor coordination, mice that failed the test were considered frail for this parameter. To calculate the “Valencia Score” of frailty, which provides an overall frailty assessment based on the five parameters, we divided the total number of failed tests by the total number of tests performed, resulting in the percentage of mice classified as frail. Statistical differences were assessed using the Chi-square test.

### Cell Culture

Human-induced pluripotent stem cells (hiPSCs) were obtained from the Gibco Human Episomal iPSC line (iPSC6.2, from WiCell Bank) ^101^ and differentiated into skeletal muscle cells following a modified, previous described protocol ^102^. Cells were maintained on a hiPSC-qualified Matrix (BD Matrigel®, BD Biosciences, NJ) in the presence of mTeSR Plus medium (Stem Cell Technologies, Vancouver, Canada) in 10 cm on Matrigel (Corning, NY) coated dishes at 37 °C with 5% O_2_ and 5% CO_2_. When the colony size reached >600 μm (0.6 mm) in diameter and the colony density on the plate was approximately 30%–40%, differentiation was induced by switching the culture medium to a myogenic differentiation medium composed of a chemically defined, serum-free medium (DMEM/F-12) supplemented with 1% (v/v) Insulin-Transferrin-Selenium-Ethanolamine and 1% (v/v) penicillin-streptomycin-glutamine (Thermo Fisher Scientific, Waltham, MA).

Starting at day 0 of differentiation, cells were cultured in the presence of 3.5 μM CHIR 99021 (Cayman, Sapphire Bioscience, Redfern, NSW, Australia) for 5 days. The culture medium was then replaced with myogenic differentiation medium supplemented with 20 ng/ml of FGF2 (PeproTech - Thermo Fisher Scientific, Waltham, MA) for 14 days. Following this, cells were cultured for an additional 16 days in myogenic differentiation medium (refreshed daily). After 35 days of differentiation, cells were purified by fluorescence-activated cell sorting. Cells were detached using 0.05% (v/v) TrypLe (Thermo Fisher Scientific, Waltham, MA), washed, and incubated with fluorochrome-labelled antibodies at a concentration of 10^6^ cells/ml for 30 min on ice (anti-HNK-1-FITC, 1:100, Aviva Systems Biology, San Diego, CA; anti-c-MET-APC, 1:50, R&D Systems, Minneapolis, MN; PE, 1μL per 10^6^ cells). After staining, cells were washed, resuspended in myogenic progenitor proliferation medium [DMEM high glucose without pyruvate (Thermo Fisher Scientific, Waltham, MA), 10% (v/v) FBS (Thermo Fisher Scientific, Waltham, MA), 100 ng/ml FGF (PeproTech - Thermo Fisher Scientific, Waltham, MA), 1% (v/v) penicillin-streptomycin-glutamine (Thermo Fisher Scientific, Waltham, MA)] and filtered through a 70 μm strainer. Live/c-MET^+^ cells were sorted using a 100 μm nozzle and collected in ice-cold myogenic progenitor proliferation supplemented with 1x RevitaCell Supplement (Thermo Fisher Scientific, Waltham, MA). The sorted cells were plated on Matrigel-coated dishes in myogenic progenitor proliferation medium, which was refreshed daily, and maintained at 37 °C with 5% O_2_ and 5% CO_2_ until they reached 90% confluence.

The differentiation of hiPSCs into myoblasts was assessed by immunofluorescence microscopy staining for MyoD. Briefly, cells were fixed with 4% (w/v) paraformaldehyde for 10 min, washed thrice, permeabilised using 0.5% (w/v) Triton X-100 in PBS for 5 min and blocked for 30 min in 10% (v/v) goat serum and 5% (w/v) bovine serum albumin in PBS. MyoD was stained with rabbit α-MyoD (Alfagene Bioscience, Farfiled, NJ; Cat # PA539265) in blocking solution supplemented with 0.1% (w/v) saponin at 4°C overnight. Cells were then washed thrice with PBS and incubated with doat *a*-rabbit IgG [H+L] Alexa Fluor 647 (Cat # A21245, Thermo Fisher Scientific, Waltham, MA), Phalloidin-iFluor (Cat # ab176756, Abcam, San Francisco, CA) and DAPI in blocking solution plus 0.1% (w/v) saponin for 1 h, before washing and mounting in Fluoromount-G™ (Cat # 00-4958-02, Thermo Fisher Scientific, Waltham, MA). Images were acquired using a a Leica Widefield Fluorescence Microscope.

### CRISPR-Cas9 gene editing

*NOX4* in human iPSC-derived myoblasts was deleted CRISPR-Cas9 ribonucleoprotein (RNP) gene editing. Cells were electroporated with recombinant Cas9 (74 pmol Alt-R S.p. Cas9 Nuclease V3; Integrated DNA Technologies, Coralville, IA) precomplexed with short guide RNAs (sgRNAs; 600 pmol; Synthego, Menio Park, CA) targeting *NOX4* (GAGGUUAAGAACAGAUGCUG) using the P3 Primary Cell 4D-Nucleofector X Kit (Lonza, Basel, Switzerland) according to the manufacturer’s instructions. After electroporation, cells were rested in supplemented DMEM cell media for recovery for 72 h before expansion and downstream analyses.

### Proteomic analysis of human mitochondrial proteins

#### Mitochondria isolation

Mitochondria were isolated from frozen skeletal muscle ^103^. Samples were mechanically fragmented using a scalpel in a Petri dish, then subjected to trypsin digestion for 30 minutes at 4°C (0.05% Trypsin, 10 mM EDTA in PBS, pH 7.4). The homogenisation process was carried out in A medium (0.32M sucrose, 1mM EDTA, 10 mM Tris-HCl, pH 7.4) using a glass tissue grinder with a motor-driven Teflon pestle (Heidolph RZR 2041), applying 20 strokes at 600 rpm. Samples were initially centrifuged at 1,000 × g for 5 minutes at 4°C, with the resulting supernatant subjected to a second centrifugation at 10,000 × g for 10 minutes at 4°C. The mitochondrial pellets were resuspended in A medium.

Mitochondria-enriched muscle samples underwent protein extraction using CS buffer (Pipes pH 6.8, MgCl2, NaCl, EDTA, sucrose, SDS, sodium orthovanadate; Biochain Institute, Inc. #K3013010-5), freshly supplemented with protease and phosphatase inhibitors. Approximately 100 μg of protein was processed for in-filter reduction and alkylation with iodoacetamide, followed by overnight trypsin digestion (Nanosep Centrifugal Devices with Omega Membrane-10K, PALL). The resulting tryptic peptides were desalted using Oasis HLB cartridges (Waters). Peptide concentration was determined using the Direct Detect infrared (IR)-based quantification system (Merck KGaA, Darmstadt, Germany). Extracted proteins (∼200 mg) underwent in-filter reduction and alkylation with iodoacetamide, followed by trypsin digestion (Nanosep Centrifugal Devices with Omega Membrane-10K, PALL). The resultant peptides were TMT-labelled according to the manufacturer’s instructions.

#### Liquid Chromatography-Tandem Mass Spectrometry (LC-MS/MS)

TMT-labelled peptides were loaded onto Evotips and washed before chromatographic separation using an Evosep One HPLC system (Endurance Column 15 cm × 150 µm ID, 1.9 µm beads-EV1106, Evosep) with a 30 SPD pre-programmed gradient. The system was coupled to a stainless-steel emitter (30 µm ID). Peptides were ionised in real time and analysed using an Orbitrap Eclipse Tribrid mass spectrometer (Thermo Fisher, San José, CA, USA) with a 2-second TopSpeed method. MS spectra were acquired in the Orbitrap analyser within a 390-1700 m/z range at a 60,000 FT resolution. HCD fragmentation was performed at 33 eV normalised collision energy, and MS/MS spectra were analysed at 30,000 resolution in the Orbitrap. A dynamic exclusion of 20 seconds was applied.

#### Protein identification and quantification

For peptide identification, MS/MS spectra were analysed using the SEQUEST HT algorithm within Proteome Discoverer 2.1^104^) (Thermo Scientific) against the Swiss-Prot database ^105^ containing mouse protein sequences (mouse_202105_sw.target-decoy.fasta, 185,234 sequences) concatenated with decoy sequences generated via DecoyPyrat ^106^. Trypsin digestion was set to allow a maximum of two missed cleavages. Fixed modifications included cysteine carbamidomethylation (57.021464 Da) and TMT labelling at the N-terminal and lysine residues (304.2071 Da), while methionine oxidation (15.994915 Da) was considered a dynamic modification. The precursor mass tolerance was set at 800 ppm, fragment mass tolerance at 0.03 Da, and precursor charge states ranged from 2-4.

False discovery rate (FDR) was determined using the corrected Xcorr score (cXcorr) ^107^ and the target/decoy competition strategy, applying the picked FDR method at the peptide level ^108^. An additional filter for precursor mass tolerance of 15 ppm was used ^109^. A 1% FDR threshold was applied for peptide identification.

Quantitative data from TMT reporter intensities were integrated at the spectrum, peptide, and protein levels using the WSPP model ^110^and the Generic Integration Algorithm (GIA) in the iSanXoT program ^111^. Protein quantitative values were expressed as the standardised variable Zq (normalised log2-ratios expressed in standard deviation units based on estimated variances).

### RNA sequencing

Total RNA was isolated from homogenised mouse gastrocnemius muscle tissue with RNeasy mini kits (Qiagen, Hilden, Germany) and processed for library preparation (PolyA, unstranded) and sequencing on an Illumina Novaseq PE150 by Azenta Life Sciences (Burlington, MA). Sequencing fastq files were processed using the nf-core-rnaseq (v3.10.1) pipeline as implemented in Laxy ^112^, and feature count matrices used for further analysis. Counts matrices for the Human dataset were downloaded from GSE159217. Differential expression was analysed using the Python implementation of DESeq2 (v0.4.2) ^113^, with genes with absolute logFoldChange >0.5 and FDR < 0.05 determined to be differentially expressed. Pathway overrepresentation was analysed using the enrichR wrapper within gseapy (v1.1.0)^114^. For ranked gene set enrichment analysis (GSEA), DESeq2 summary statistics for each gene were ranked by the Wald statistic, and pathway enrichment assessed with gseapy after excluding outliers (p-value = NA) and any duplicated gene names. Heatmaps were generated using the log of the normalised counts generated by DESeq2.

### Real Time PCR

Total RNA was extracted using RNAzol (Sigma-Aldrich, St. Louis, MO), and its quality and quantity were assessed using a NanoDrop 3300 spectrophotometer (Thermo Fisher Scientific, Waltham, MA). Complementary DNA (cDNA) was synthesized from mRNA using the High-Capacity cDNA Reverse Transcription Kit (Applied Biosystems, Foster City, CA). Quantitative real-time PCR (qRT-PCR) was performed using either the TaqMan Universal PCR Master Mix with TaqMan Gene Expression probes (Applied Biosystems) or the Quantinova SYBR Green Master Mix (Qiagen, Hilden, Germany) with either Bio-Rad Prime PCR primers or validated oligonucleotide primers. Housekeeping genes *Gapdh or Rn18s* were used as internal controls for tissue samples, while *Rplp0* was used for cell samples. *Rn18s* was utilised for the aging studies. Thermal cycling and fluorescence signal detection were conducted using the QuantStudio™ 5 (Thermo Fisher Scientific, Waltham, MA). Relative gene expression levels were determined using the ΔΔCT method. All primers utilised are described below:

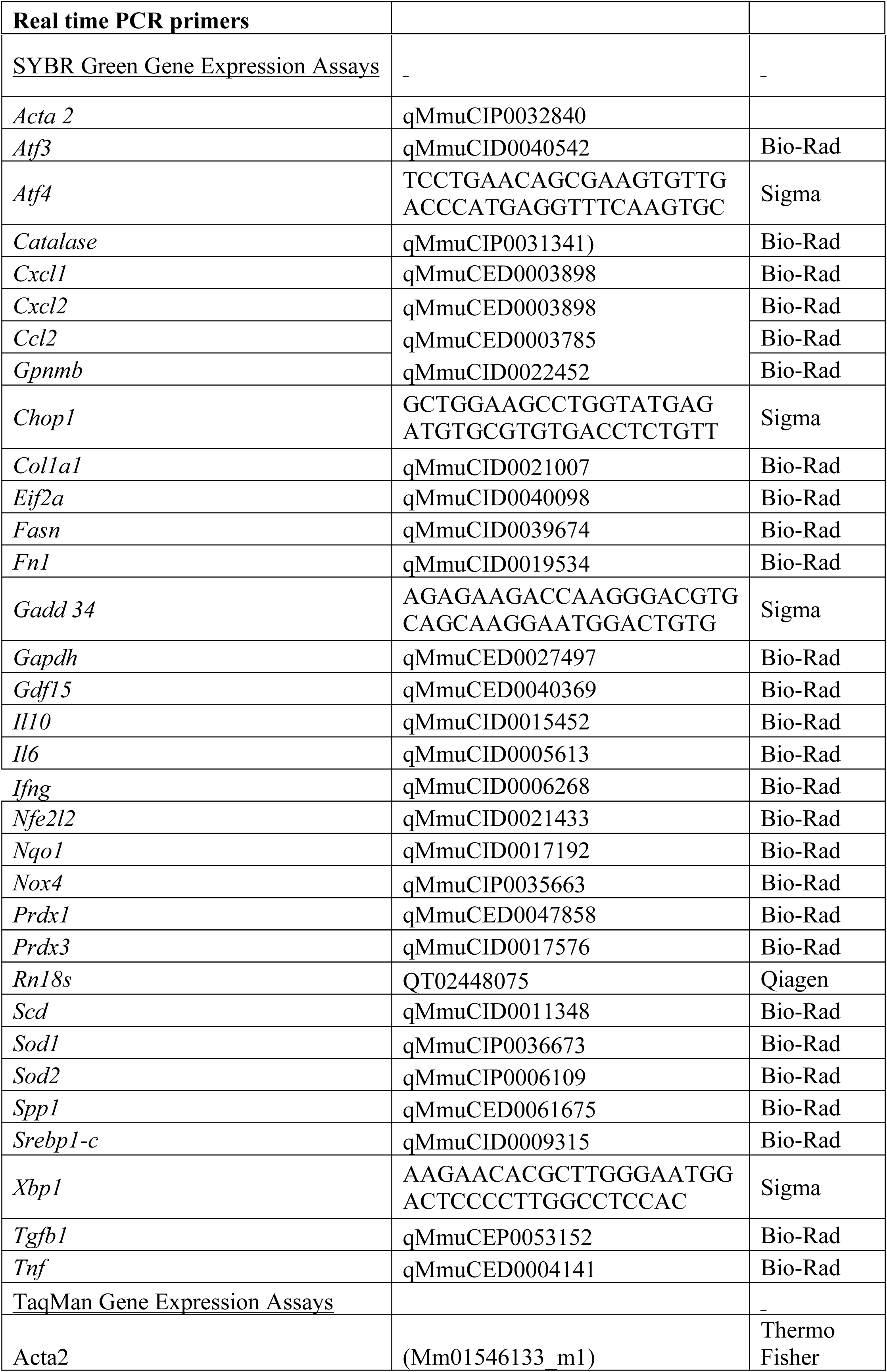
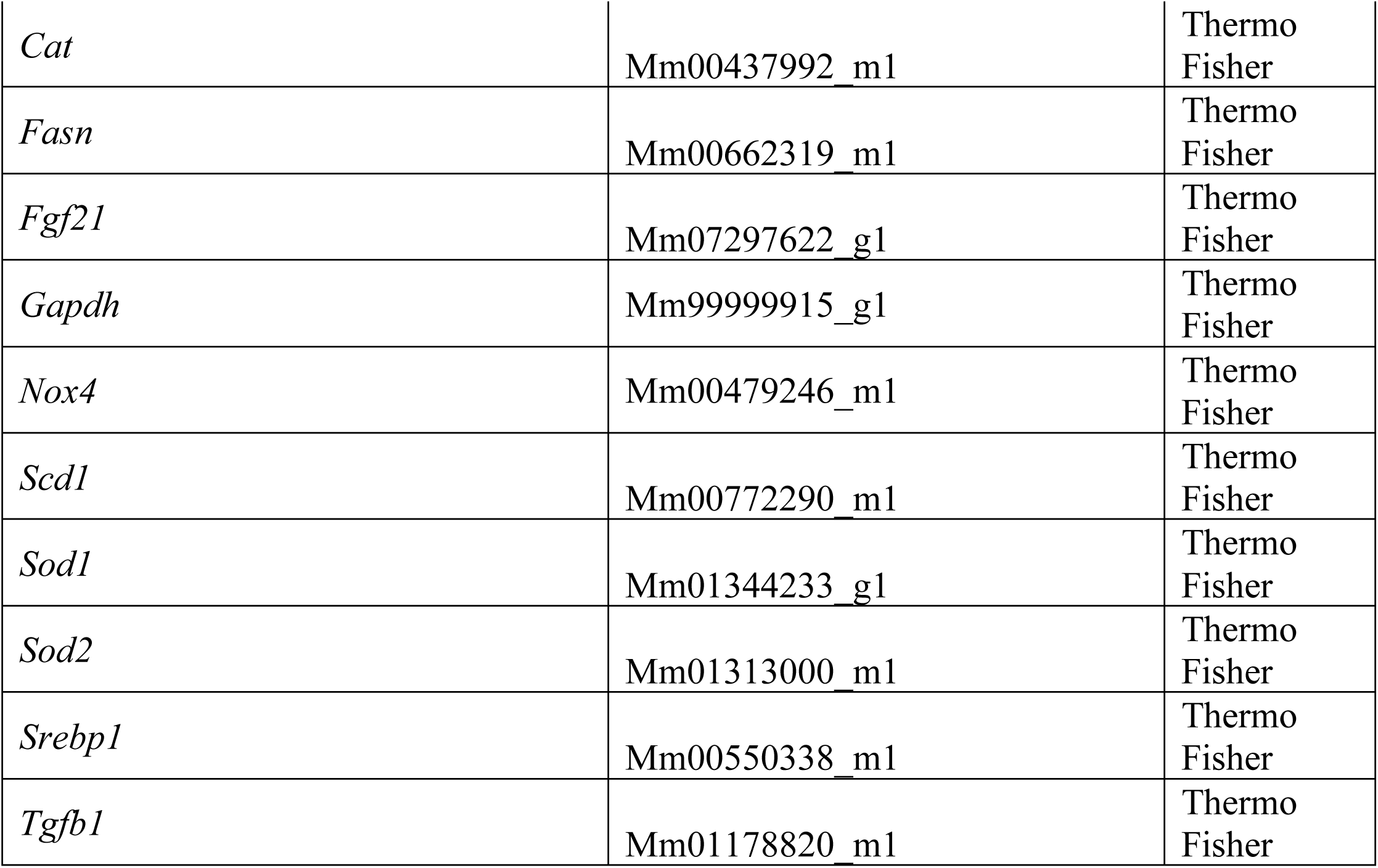

### Immunoblotting

Murine or human tissues snap-frozen in liquid nitrogen were homogenized using a bead homogenizer (Bead Ruptor 12, Omni International, GA) with 1 mm diameter zirconia/silica beads (Biospec Products, OK) for 30–60 s in 10–20 volumes of ice-cold RIPA lysis buffer (50 mM HEPES [pH 7.4], 1% (v/v) Triton X-100, 1% (v/v) sodium deoxycholate, 0.1% (v/v) SDS, 150 mM NaCl, 10% (v/v) glycerol, 1.5 mM MgCl2, 1 mM EGTA, 50 mM NaF, 5 µg/ml leupeptin, 1 µg/ml pepstatin A, 1 mM benzamidine, 2 mM phenylmethylsulfonyl fluoride, 1 mM sodium orthovanadate). Homogenates were incubated on ice for 30 min and clarified by centrifugation at 16,000 g for 30 min at 4°C. Clarified tissue lysates were resolved by SDS-PAGE and transferred onto PVDF membranes for immunoblotting.

### Skeletal muscle histology and immunostaining

Gastrocnemius muscles were dissected from 6-, 12- and 20-month old mice, snap frozen in liquid nitrogen cooled isopentane and stored at -80°C until required. Transverse muscle cryosections (10 μm) were prepared and used for all histology and immunostaining experiments as described previously ^28^.

Haemotoxylin and eosin (H&E), and succinate dehydrogenase (SDH) staining were viewed via slide scanning using either an Olympus DotSlide microscope where individual images were stitched together using the VS-ASW program (Olympus) to generate a composite image of the entire muscle section, or using an Aperio Scanscope AT Turbo slide scanner (Leica Biosystems). TOMM20 immunostaining of mitochondria or fibre-type staining in muscle sections were imaged using a Leica SP5 5-Channel confocal microscope. All microscopy was performed at Monash MicroImaging or the Monash Histology Platform, Monash University, Australia. Automated fibre-typing analysis was performed using the MuscleJ2 plugin installed in ImageJ 1.54e (National Institutes of Health, Bethesda, MD, USA).

## QUANTIFICATION AND STATISTICAL ANALYSIS

Data are presented as mean ± standard error of the mean (SEM). Statistical significance was determined using two-tailed Student’s t-tests for pairwise comparisons or one-way/two-way ANOVA with multiple comparisons for group-based analyses. When there was no normal distribution at the data as assessed by Kolmogorov-Smirnov normality test, Mann-Whittney test was performed (check proper name). A p-value threshold of *p* < 0.05 was considered statistically significant (*p* < 0.05, p < 0.01, *p < 0.001, **p < 0.0001). Sample sizes and detailed statistical parameters for each experiment are indicated in figure legends.

