## Supplementary material for "A decline in skeletal muscle NOX4 abrogates adaptive homeostasis and exacerbates ageing": Supp Figure 1-16

***Table S1. NFE2L2 gene set used for enrichment analyses***

| Mouse | Human |
|---|---|
| *Cat* | *CAT* |
| *G6pdx* | *G6PD* |
| *Gclc* | *GCLC* |
| *Gclm* | *GCLM* |
| *Gpx1* | *GPX1* |
| *Gpx3* | *GPX3* |
| *Gpx4* | *GPX4* |
| *Gpx7* | *GPX7* |
| *Gsr* | *GSR* |
| *Idh1* | *IDH1* |
| *Me1* | *ME1* |
| *Nfe2l2* | *NFE2L2* |
| *Nqo1* | *NQO1* |
| *Nqo2* | *NQO2* |
| *Phgdh* | *PHGDH* |
| *Prdx1* | *PRDX1* |
| *Prdx2* | *PRDX2* |
| *Prdx3* | *PRDX3* |
| *Sod1* | *SOD1* |
| *Sod2* | *SOD2* |
| *Txn* | *TXN* |
| *Txnrd1* | *TXNRD1* |

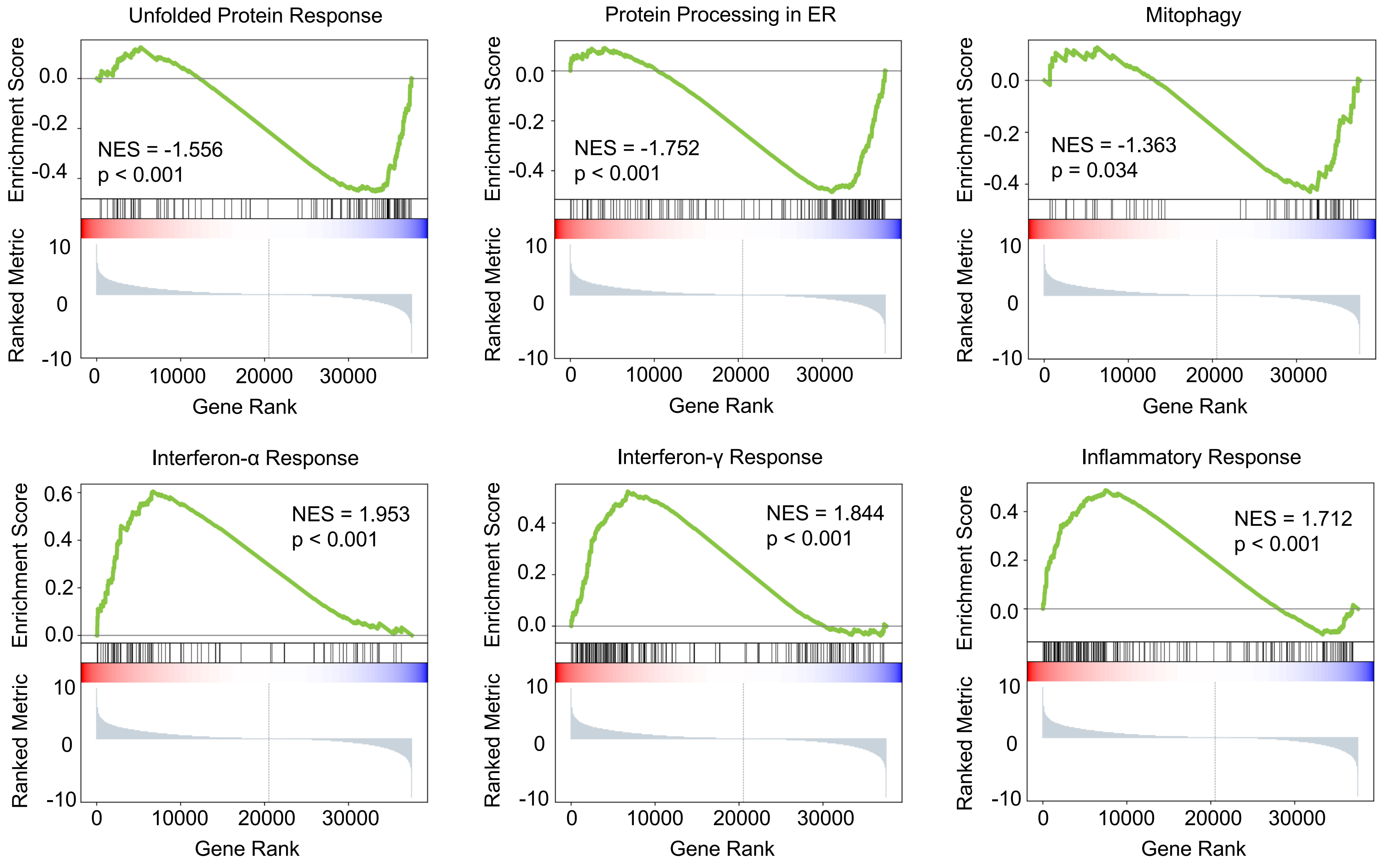


Related to Figure 1 A-C

**Figure S1**

***Figure S1. Barcode plots from ranked GSEA of RNAseq analysis of vastus lateralis in ageing human males - Related to Fig. 1 a-c.*** Barcode plots demonstrating negative enrichment of Unfolded Protein Response (Hallmark gene sets) and ER processing and Mitophagy (KEGG Pathways) in older males, and positive enrichment of Interferon α & γ Response and Inflammatory Response (Hallmark gene sets).

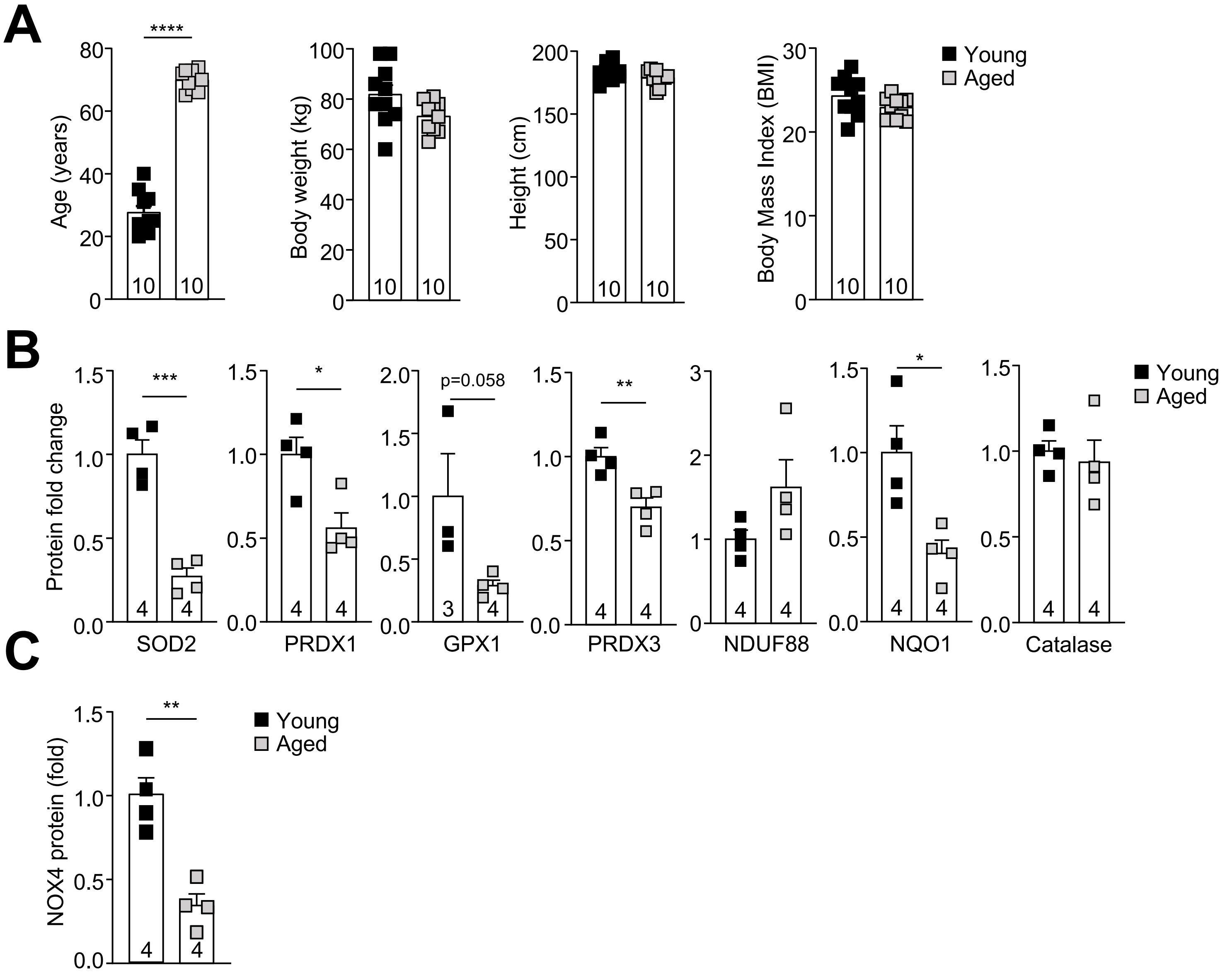


Related to Figure 1D-F

Figure S2

***Figure S2. Antioxidant defence proteins decline in vastus lateralis muscles of aged men- Related to Fig. 1d, g-h.*** Skeletal muscle *vastus lateralis* biopsies from young (27.6 ± 6.6 years old) and aged (69.9 ± 3.1 years old) men wer used for analysis of antioxidant defence protein abundance. Anthropometric characteristics of the participants (**a**) including age, body weight, height and BMI. **b-c**) Biopsies were processed for immunoblotting to assess the abundance of  antioxidant enzymes and NOX4; quantified results of results in b) Figure 1d and c) Figure 1h are shown. Results are mean ± SEM for the indicated number of male participants; significance determined by Student's t-test.

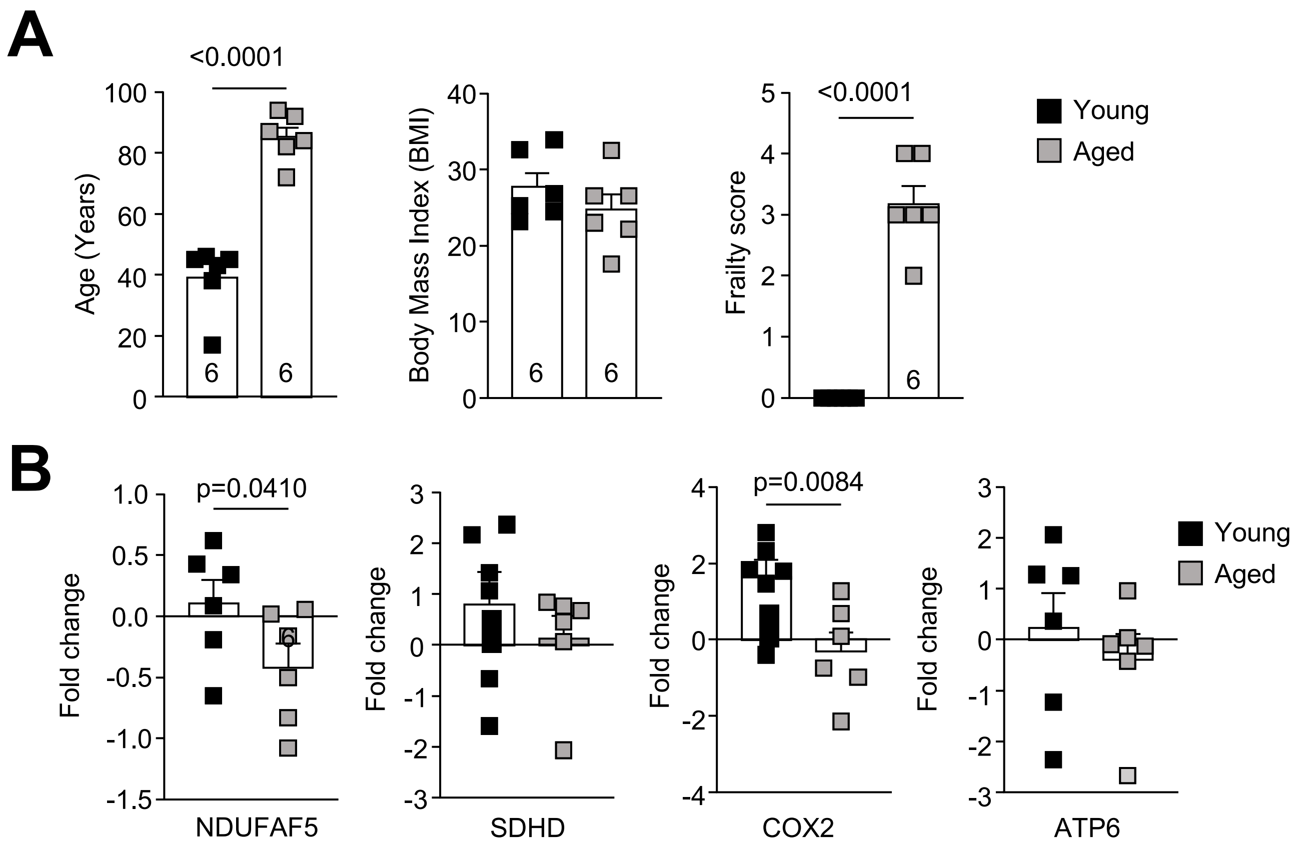


Related to Figure 1G-H

**Figure S3**

***Figure S3. Proteomics analyses of human skeletal muscle mitochondria-enriched fractions** - **Related to Fig. 1e-f.*** Mass spectrometry-based proteomic analyses of mitochondria-enriched fractions from hip skeletal muscle biopsies of BMI-matched aged, physically inactive (86.7 ± 9.7 year old) versus young, physically fit (37.3 ± 10.6 years old) individuals. (**a**) Anthropometric characteristics of participants including age, BMI, Valencia Frailty score. **b**) Mitochondria-enriched fractions were analysed for the abundance of oxidative phosphorylation complexes. Representative and quantified results are shown (mean ± SEM) for the indicated number of human mitochondrial fractions; significance determined by Student's t-test.

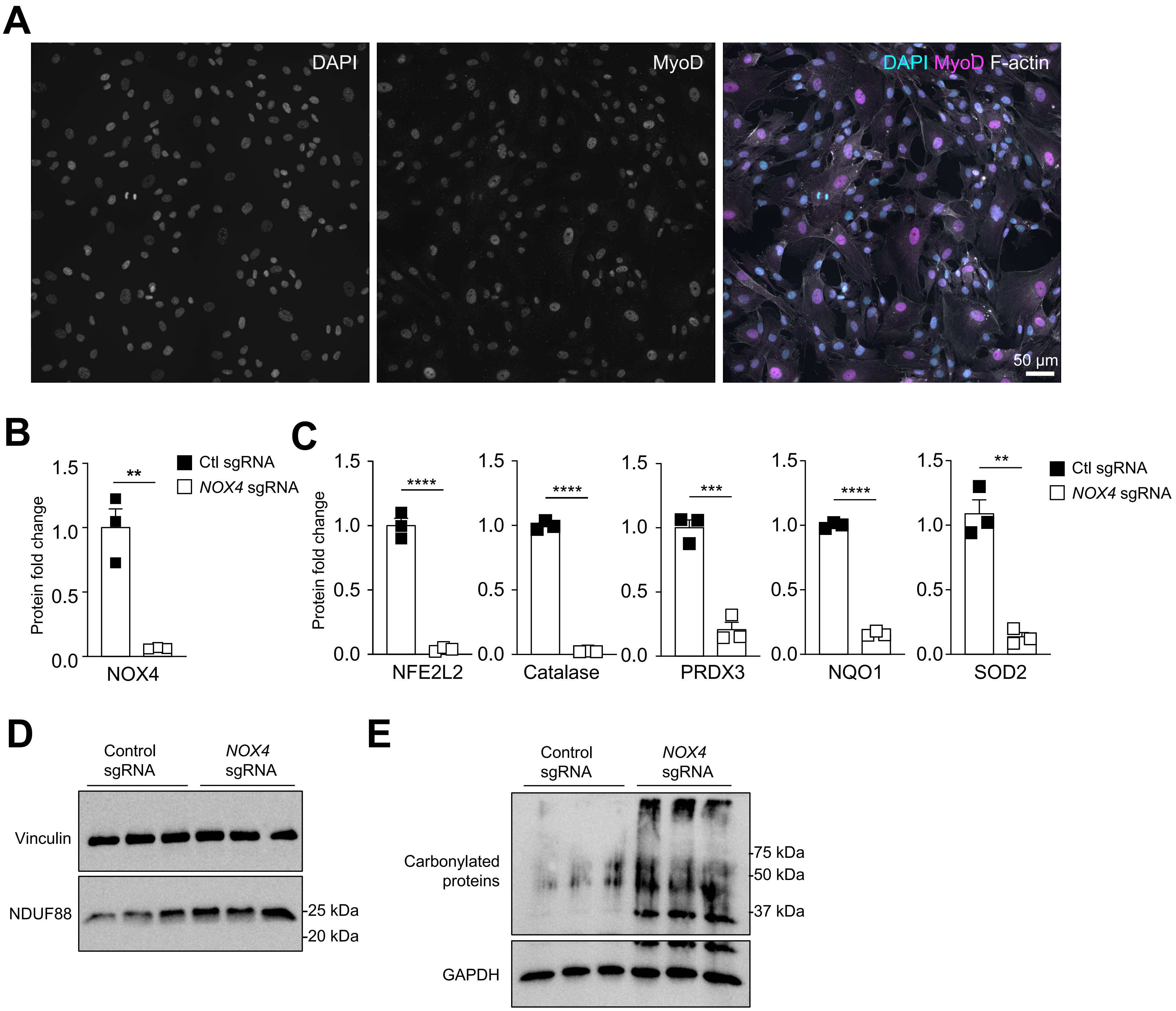


Related to Figure 1 I

**Figure S4**

***Figure S4. NOX4 deletion in human iPSC-derived myoblasts. - related to Fig. 1l.*** Human induced pluripotent stem cells (iPSCs) were differentiated into myoblasts. **a**) Differentiation into myoblasts was validated by the expression of MyoD assessed by immunofluorescence microscopy staining for MyoD, DAPI and F-actin. **b-d**) *NOX4* was deleted in iPSC-derived myoblasts by CRISPR RNP gene-editing using control or *NOX4* specific sgRNAs and the abundance of **b**) NOX4 protein and **c**) antioxidant defence proteins assessed by immunoblotting; quantified results of b-c) Figure 1l are shown. Alternatively control and NOX4-deficient iPSC-derived human myoblsts were processed for immunoblotting monitoring for **d**) the abundance of the mitochondrial complex protein NDUF88, or **e**) protein carbonylation. Representative and quantified results are shown (means ± SEM) for the indicated number of experiments; significance determined by Student's t-test (b-c).

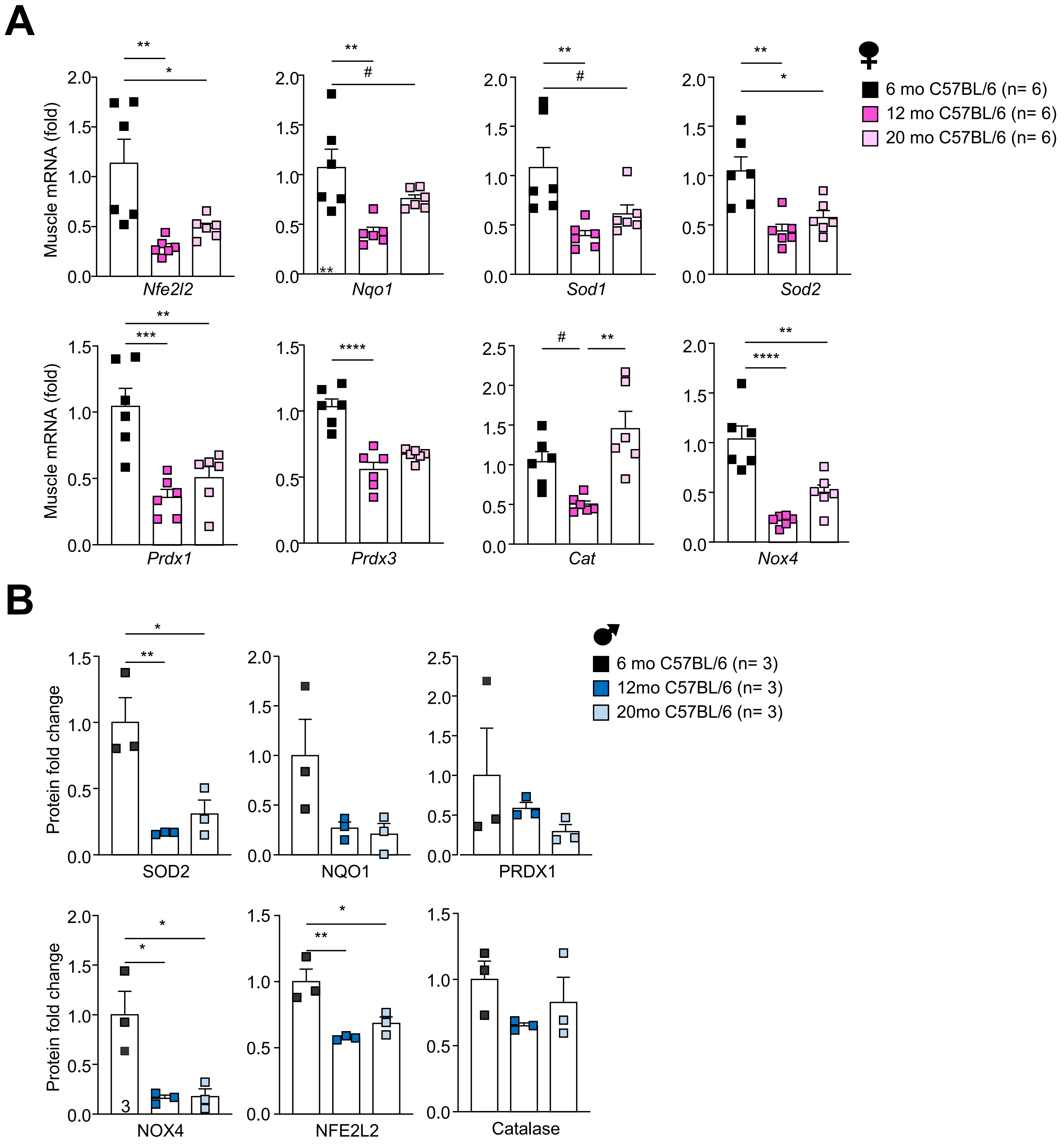


Related to Figure 2

**Figure S5**

***Figure S5. Skeletal muscle antioxidant defence declines during ageing in C57BL/6 mice - Related to Fig 2a-b.*** **a**) Female C57BL/6 mice were fed a standard chow-diet (4.8% fat) for 6, 12 or 20 months and *gastrocnemius* muscles extracted and analysed by qPCR. **b**) Male C57BL/6 mice were fed a standard chow diet (4.8% fat) for 6, 12 or 20 months and gastrocnemius muscles extracted and processed immunoblotting; quantified results of Figure 2c are shown. Results shown are means ± SEM for the indicated number of mice; significance determined using one-way ANOVA or where indicated (#) Student's t-test.

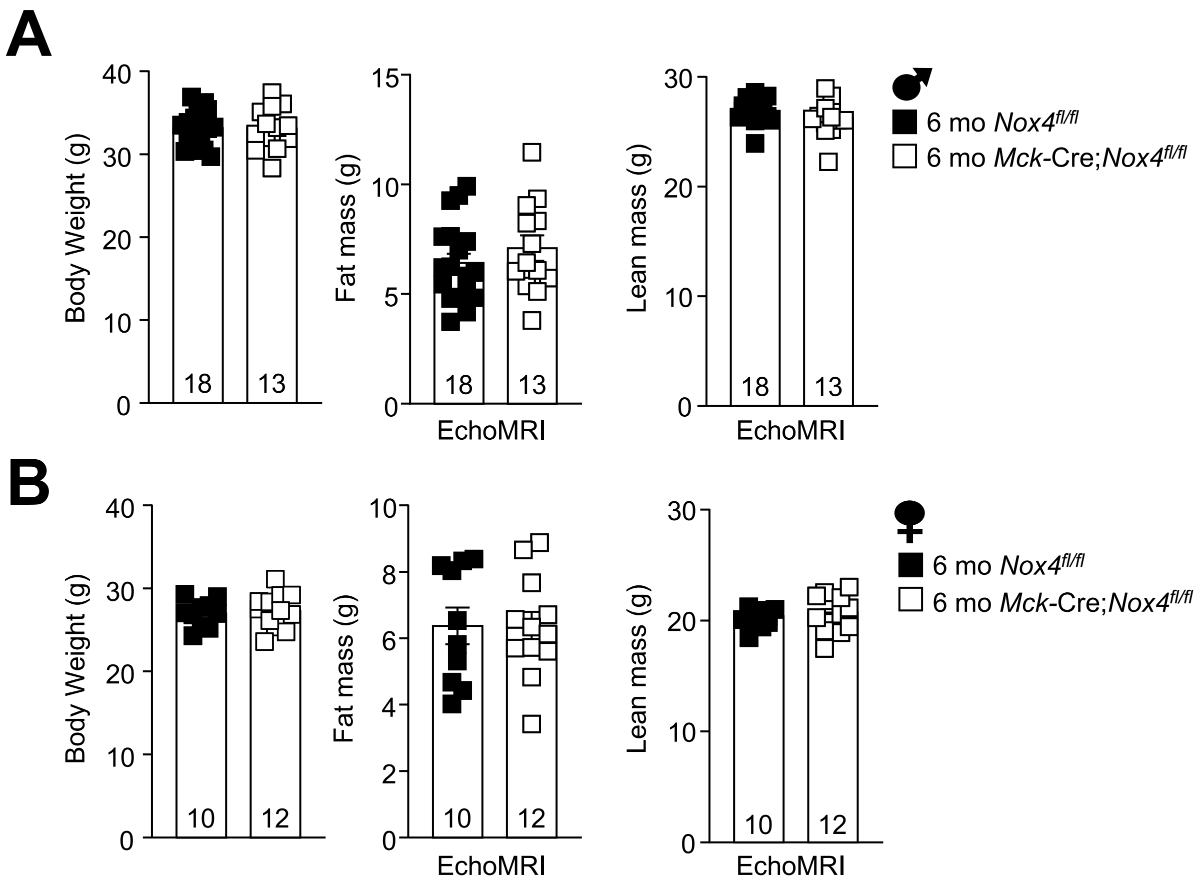


Related to Figure 3A

***Figure S6. Muscle NOX4-deficiency does not alter body weight or body composition at 6 months of age - Related to Fig 3a-d.*** Body weights and body composition (EchoMRI) in 6 month old *Nox4*$^{fl/fl}$ and *Mck*-Cre;*Nox4*$^{fl/fl}$ **a**) male and **b**) female mice fed a standard chow diet (4.8% fat). Results shown are means ± SEM for the indicated number of mice.

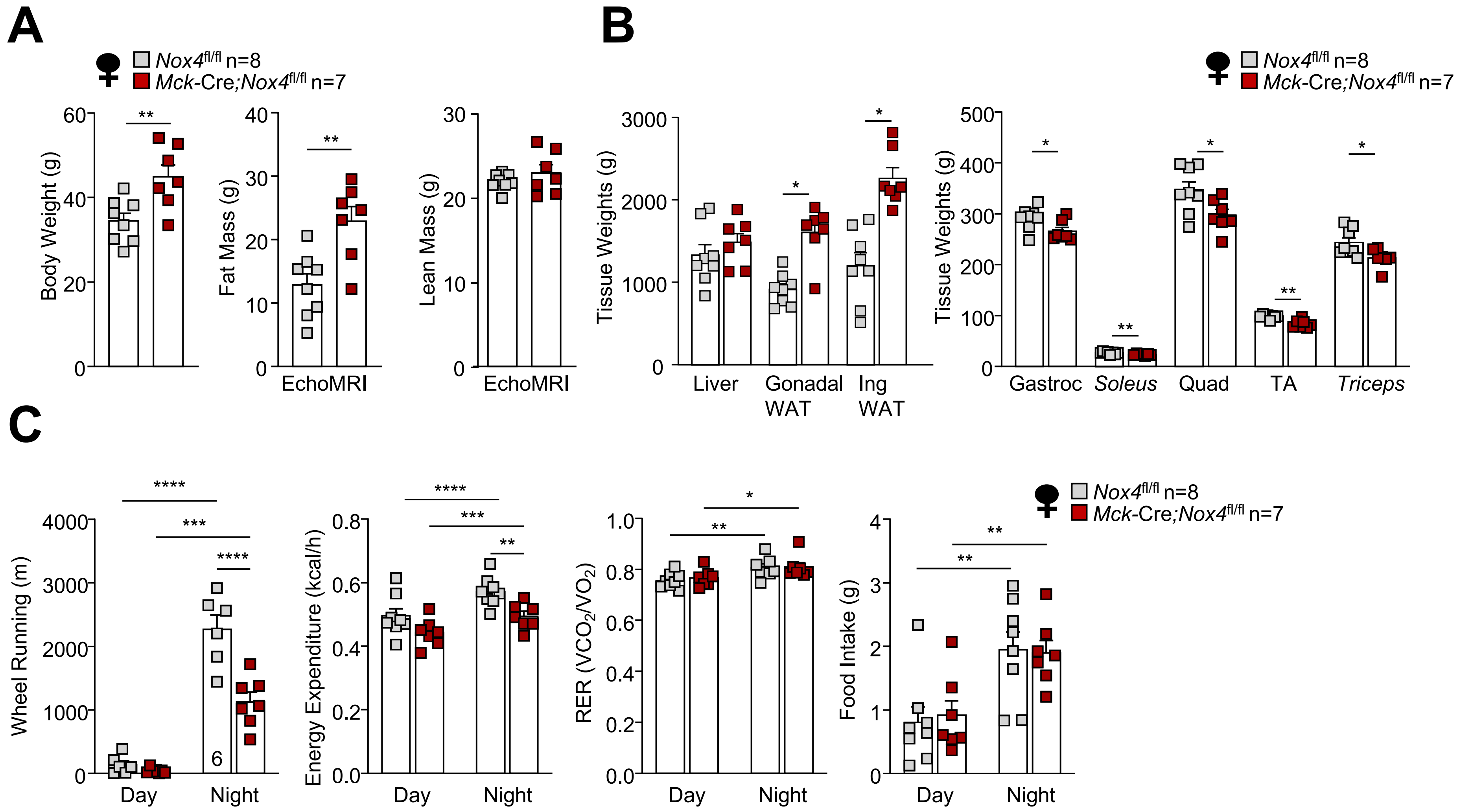
A
B
C
Nox4fl/fl n=8
Mck-Cre;Nox4fl/fl n=7
Body Weight (g)
Fat Mass (g)
Lean Mass (g)
EchoMRI
Tissue Weights (g)
Liver
Gonadal WAT
Ing WAT
Gastroc
Soleus
Quad
TA
Triceps
Wheel Running (m)
Energy Expenditure (kcal/h)
RER (VCO2/VO2)
Food Intake (g)
Day
Night



**Figure S7**

***Figure S7. Muscle NOX4-deficiency increases adiposity and decreases muscle mass and voluntary wheel running in aged female mice – Related to Fig. 3a-d.*** *Nox4*$^{fl/fl}$ and *Mck*-Cre;*Nox4*$^{fl/fl}$ female mice were fed a standard chow diet (4.8% fat) for 20-months. **a**) Body weights and body composition (EchoMRI). **b**) Tissue weights. **c**) Ambulatory activity (wheel running), energy expenditure, RERs and food intake were analsed in metabolic cages (Promethion). Representative and quantified results are shown (means ± SEM) for the indicated number of mice; significance determined by Student's t-test (b) or two-way ANOVA (c).

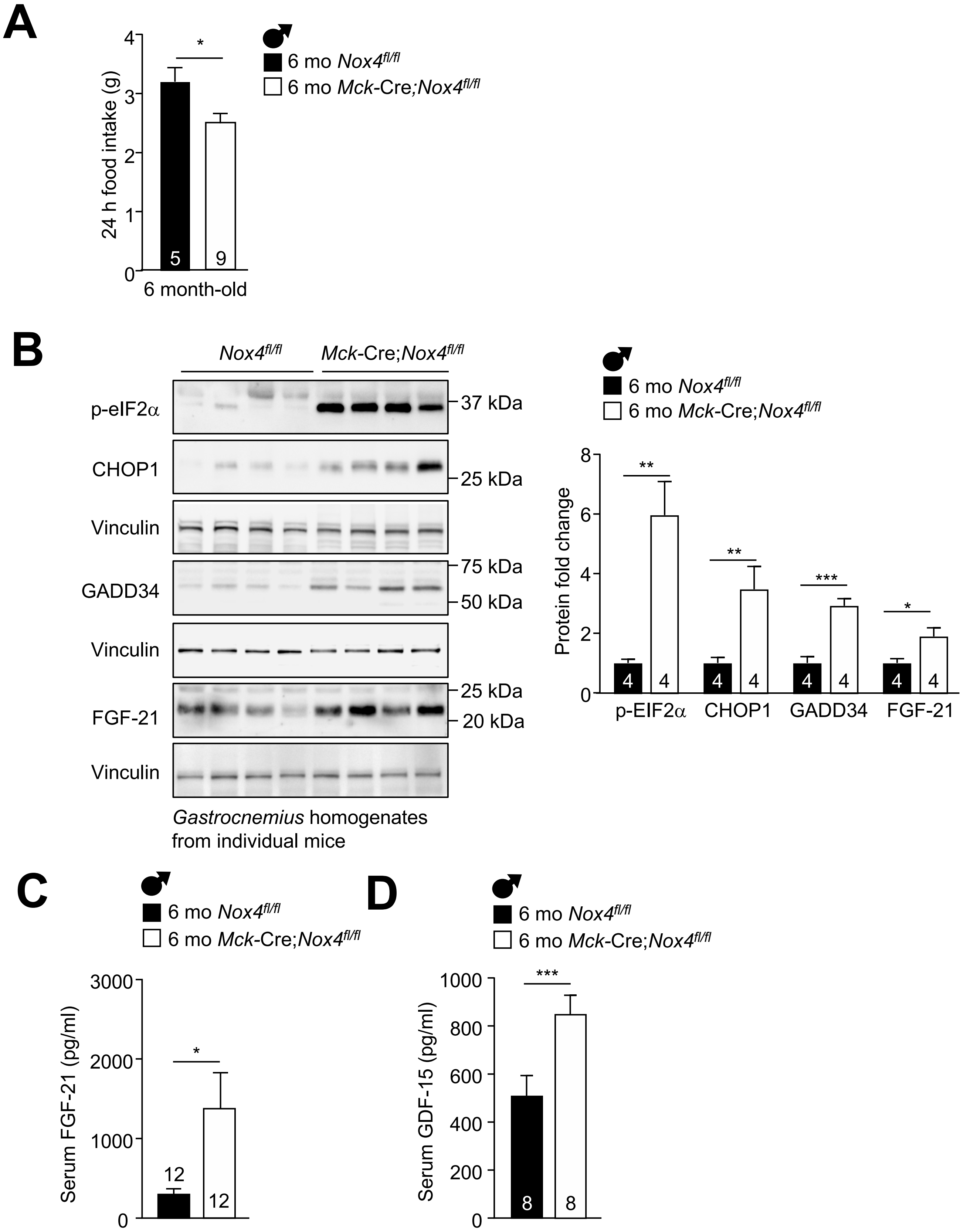


Related to Figure 3A-H

**Figure S8**

***Figure S8. Muscle NOX4-deficiency drives a robust integrated stress response in 6 month old mice - Related to Fig. 3a-d.*** *Nox4*$^{fl/fl}$ and *Mck*-Cre;*Nox4*$^{fl/fl}$ male mice were fed a standard chow diet (4.8% fat) for 6 months. **a**) 24 h food intake. **b**) *Gastrocnemius* muscles were processed for immunoblotting to assess eIF2α Ser-51 phosphorylation (p-eIF2α) and CHOP-1, GADD34 and FGF21 protein levels. **c**) Serum FGF-21 and **d**) GDF-15 levels from ad libitum chow fed mice. Representative and quantified results are shown (means ± SEM) for the indicated number of mice; significance determined by Student's t-test.

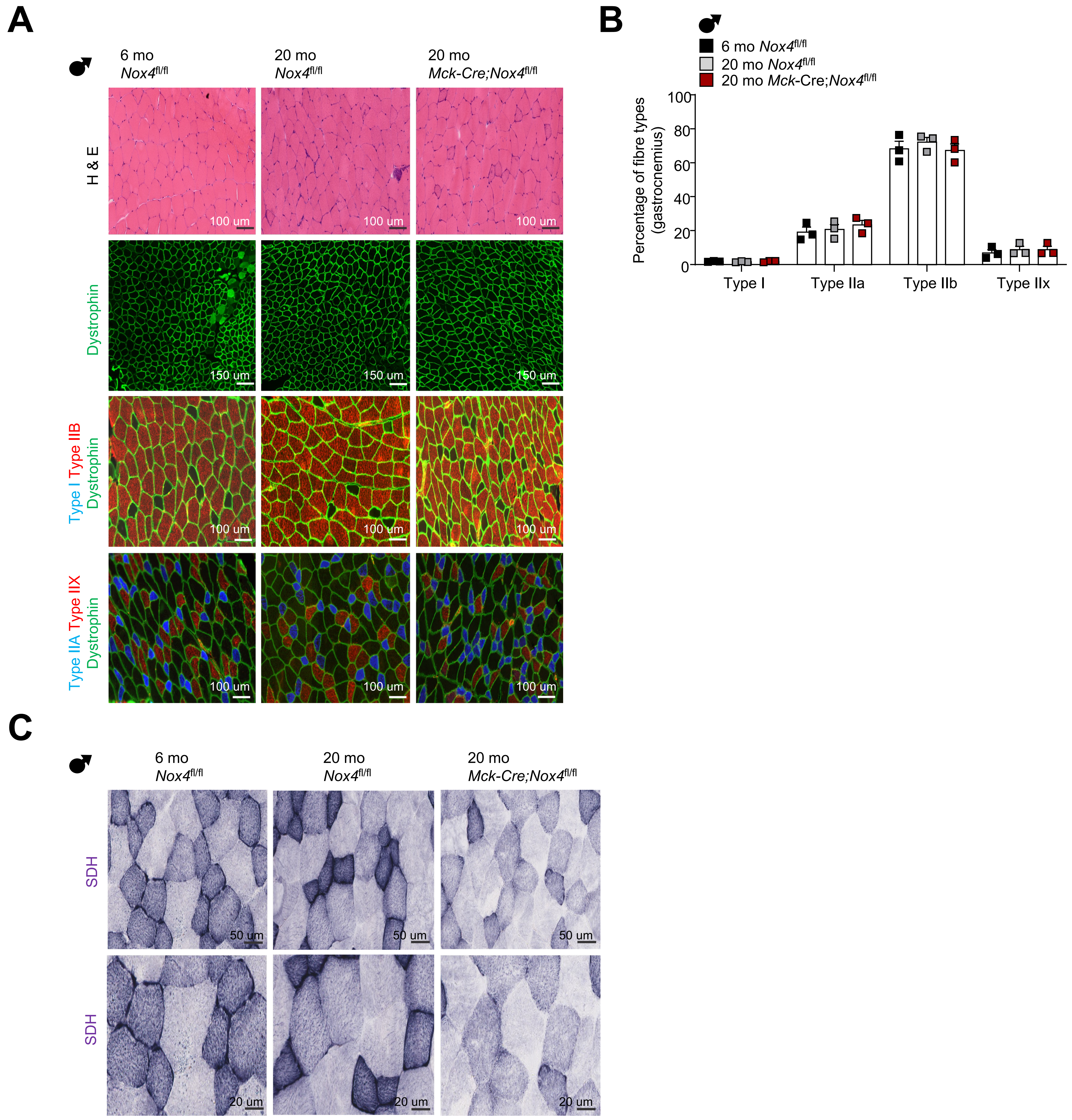


Related to Figure 3

**Figure S9**

***Figure S9. Muscle NOX4-deficiency does not alter fibre type but decreases mitochoindrial content in aged mice – Related to Fig 3f-g.*** *Nox4*$^{fl/fl}$ control and *Mck*-Cre;*Nox4*$^{fl/fl}$ male mice were fed a standard chow diet (4.8% fat) for 6 or 20 months as indicated. **a-b**) *Gastrocnemius* muscles were dissected and processed for immunostaining. Frozen transverse muscle cryosections (10 μm) were processed for a) haemotoxylin and eosin (H&E) staining and immunostained for dystrophin, or fibre types I, or IIa, IIb, IIx and b) fibre type composition determined. **c**) Alternatively, transverse sections were processed for Succinate Dehydrogenase (SDH) staining to monitor for mitochondria content. Representative and quantified results are shown (means ± SEM) for the indicated number of mice.

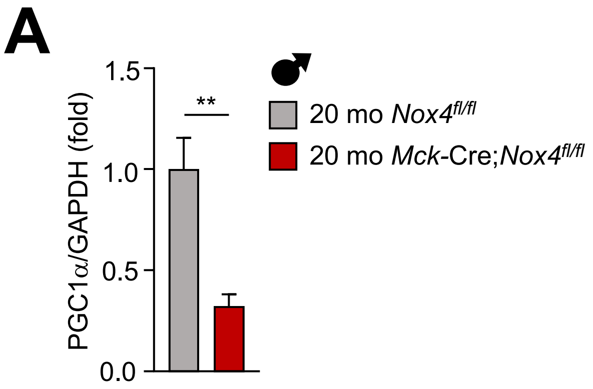


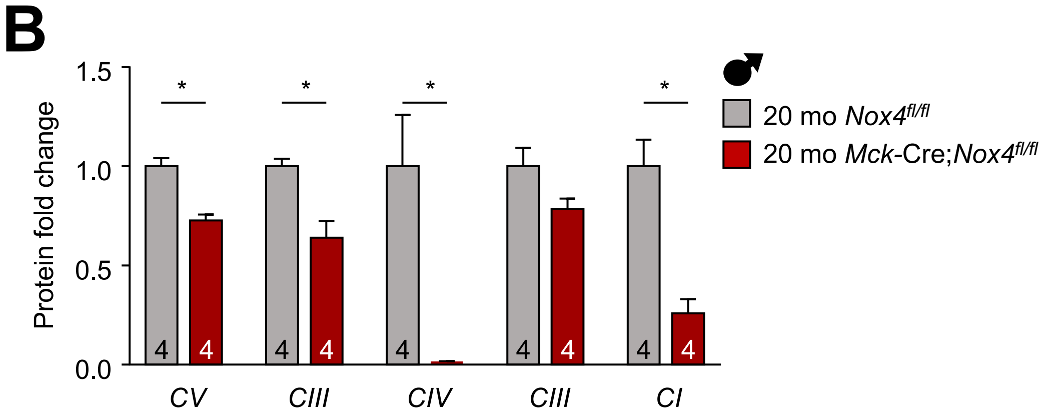


Related to Figure 3

**Figure S10**

***Figure S10 – Muscle NOX4-deficiency decreases mitochondrial content in aged mice Related to Fig. 3f***. *Nox4*$^{fl/fl}$ control and *Mck*-Cre;*Nox4*$^{fl/fl}$ muscle-specific NOX4-deficient male mice were fed a standard chow diet (4.8% fat) for 20-months. *Gastrocnemius* muscles were dissected and processed for immunoblotting to assess mmunoblotting to monitor PGC1α protein levels and OXPHOS protein complexes; quantifed results from Fig. 3f are shown. Results shown are means ± SEM for the indicated number of mice.

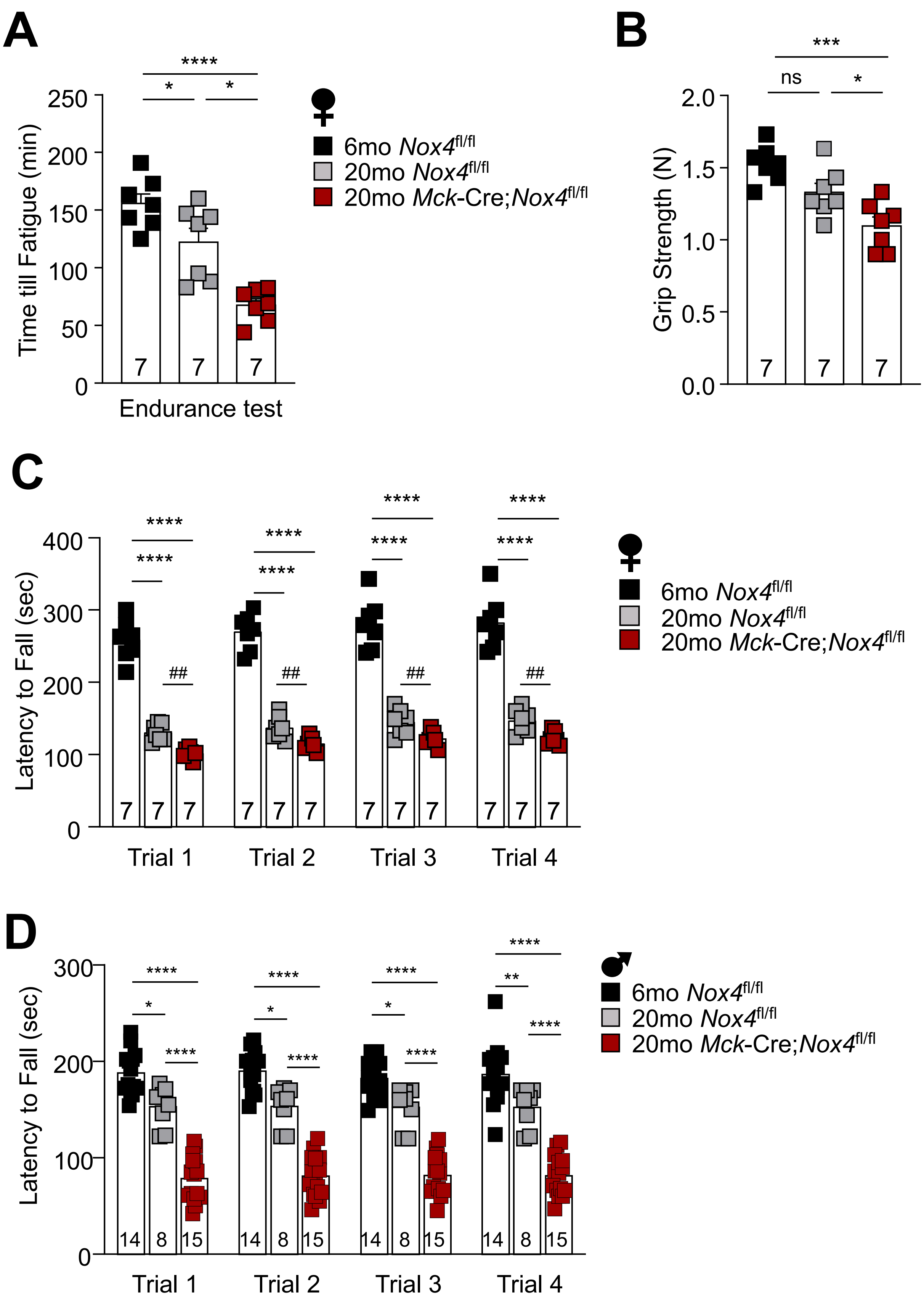


Related to Figure 3

**Figure S11**

***Figure S11. Muscle NOX4-deficiency exacerbate the decline in musle function in aged mice – Related to Fig. 3h.*** **a)** 6 or 20 month old *Nox4*$^{fl/fl}$ and Mck-Cre;*Nox4*$^{fl/fl}$ female mice as indicated were fed a standard chow diet (4.8% fat) and subjected to endurance tests, grip strength measurements, and locomotor activity (rotarod). **b**) 6 or 20 month old *Nox4*$^{fl/fl}$ and Mck-Cre;*Nox4*$^{fl/fl}$ male mice as indicated were fed a standard chow diet (4.8% fat) and subjected to rotarod tests to assess locomtor activity (trials 1-4 of results show in Fig. 3h). Results shown are means ± SEM for the indicated number of mice; significance determined using one-way ANOVA.

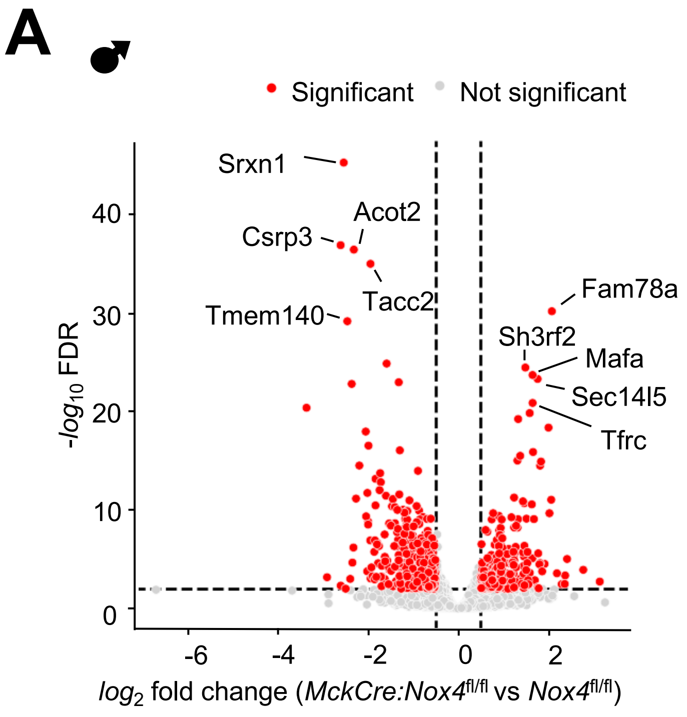


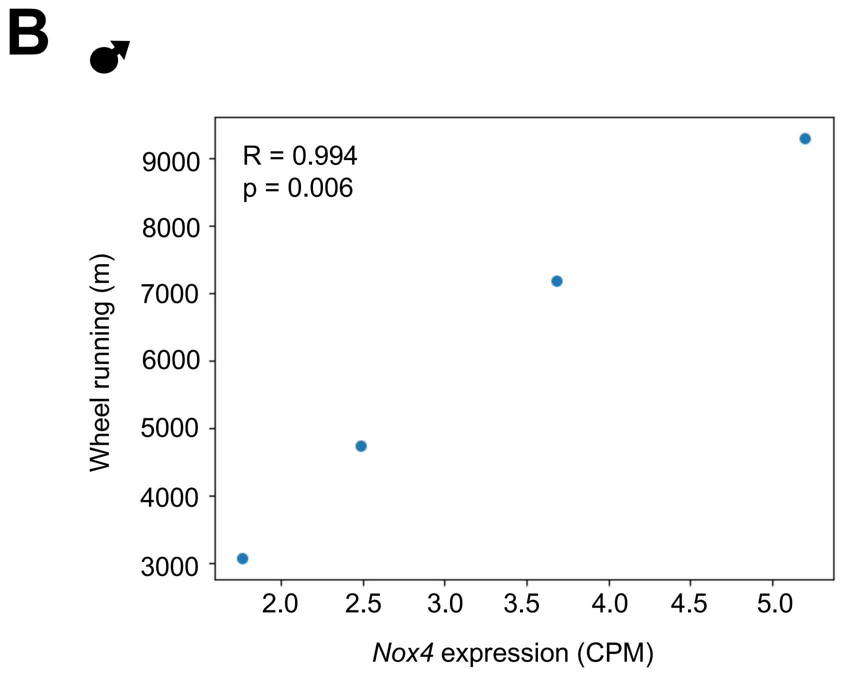


Related to Figure 4

**Figure S12**

***Figure S12. Transcriptomic analysis of muscle NOX4-deficient aged mice - Related to Fig. 4a-c.*** *Gastrocnemius* from 21 month old *Nox4*$^{fl/fl}$ and *Mck*-Cre;*Nox4*$^{fl/fl}$ male chow (4.8% fat) fed mice (n=4 per genotype) was processed for bulk RNAseq. **a**) Volcano plot of 443 genes significantly upregulated and 580 significantly downregulated (FDR <0.05, absolute log fold change > 0.5). The top 5 most significant up/downregulated genes are indicated by name. **b**) Correlation between wheel running distance (m in a 2 day/night period) and *Nox4* expression (counts per million) in 21 month-old *Nox4*$^{fl/fl}$ control mice. Near perfect correlation was observed as assessed by Pearson test (R=0.994, p=0.006).

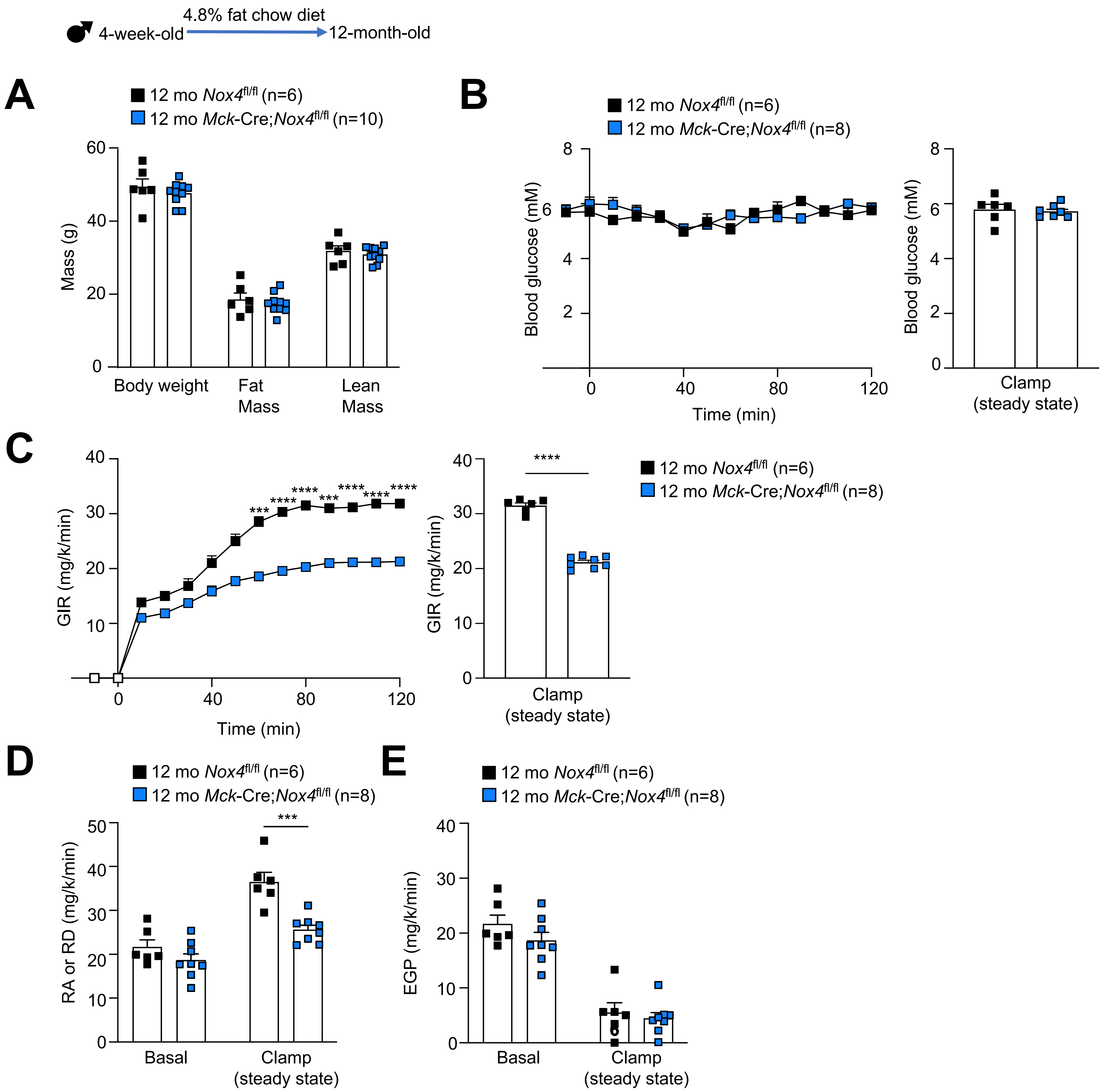


Related to Figure 5

**Figure S13**

***Figure S13. Systemic insulin resistance in 12 month old muscle NOX4-deficient mice prior to body weight changes. Related to Fig. 5c-e.*** *Nox4*$^{fl/fl}$ and *Mck*-Cre;*Nox4*$^{fl/fl}$ male mice were fed a standard chow diet (4.8% fat) for 12 months. **a**) Body weights and body composition were assessed. **b-e**) Hyperinsulinaemic-euglycemic clamps in conscious and free-moving mice. **b**) Blood glucose was measured during 120 min (line graph) as well as during the last 30 min of the clamp (80-120 min) where steady state hyperinsulinaemic-euglycaemic conditions were achieved. **c**) The glucose infusion rate (GIR), **d**) rate of glucose disappearance (RD) and **e**) endogenous glucose production (EGP) were assessed. Representative and quantified results are shown (means ± SEM) for the indicated number of mice; significance determined using a two-way ANOVA (d-e) or Student's t-test (c).

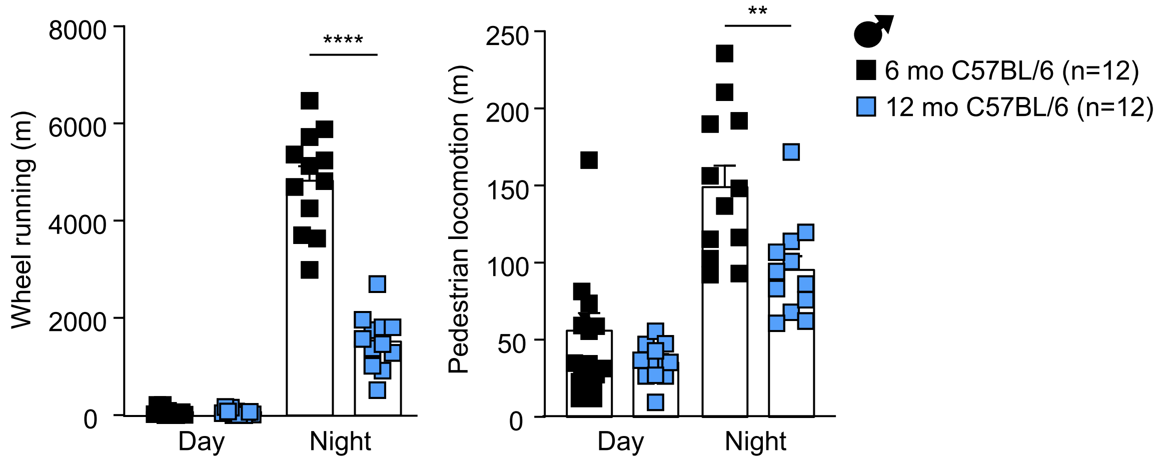


Related to Figure 6

**Figure S14**

***Figure S14. Pedestrian locomotion and wheel running decline in 12 month-old mice -Related to Fig. 6.*** C57BL/6 male mice were fed a standard chow diet (4.8% fat) for 6 or 12 months and wheel running and pedestrian locomotion assessed in metabolic cages (Promethion). Results shown (means ± SEM) for the indicated number of mice; significance determined using a two-way ANOVA.

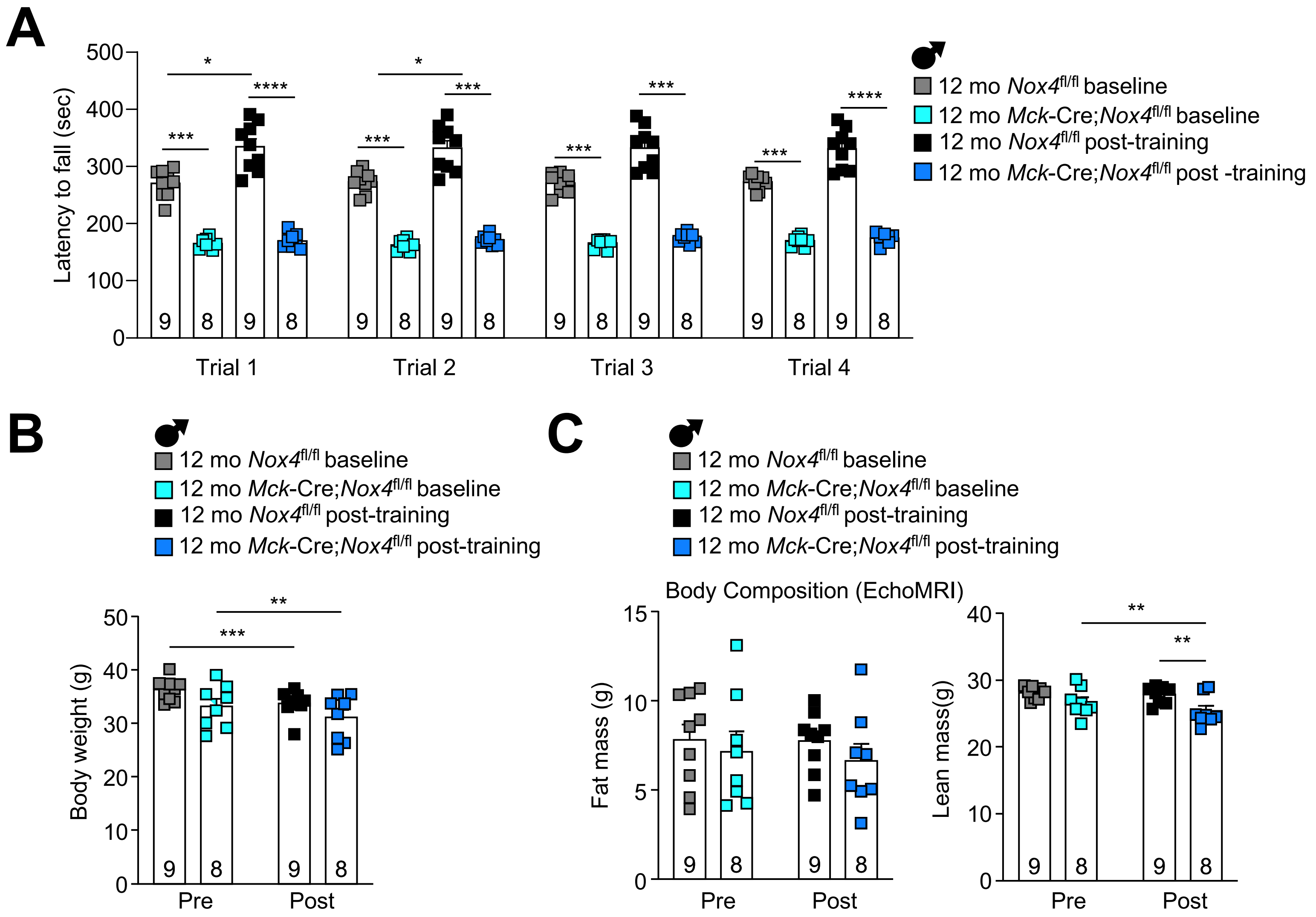
A
Latency to fall (sec)
500
400
300
200
100
0
*
***
****
*
***
***
***
***
***
****
9 8 9 8
Trial 1
Trial 2
Trial 3
Trial 4
♂
12 mo Nox4fl/fl baseline
12 mo Mck-Cre;Nox4fl/fl baseline
12 mo Nox4fl/fl post-training
12 mo Mck-Cre;Nox4fl/fl post -training
B
♂
12 mo Nox4fl/fl baseline
12 mo Mck-Cre;Nox4fl/fl baseline
12 mo Nox4fl/fl post-training
12 mo Mck-Cre;Nox4fl/fl post-training
Body weight (g)
50
40
30
20
10
0
**
***
Pre
Post
C
♂
12 mo Nox4fl/fl baseline
12 mo Mck-Cre;Nox4fl/fl baseline
12 mo Nox4fl/fl post-training
12 mo Mck-Cre;Nox4fl/fl post-training
Body Composition (EchoMRI)
Fat mass (g)
15
10
5
0
Lean mass(g)
40
30
20
10
0
**
**
Pre
Post



**Figure S15**

***Figure S15. Exercise training does not alter body weight or composition in 12 month-old muscle NOX4-deficient male mice - Related to Fig 6e-f.*** *Nox4*$^{fl/fl}$ and *Mck*-Cre;*Nox4*$^{fl/fl}$ male mice were fed a standard chow diet (4.8% fat) for 12 months and underwent pre and post exercise training (5 weeks with progressively increasing running speeds each week). **a**) Mice subjected to rotarod tests to assess locomotor activity pre and post exercise training (trials 1-4 or results shown in Fig. 6e). **b**) Body weights, **c**) body composition (EchoMRI) pre and post exercise training. Results shown means ± SEM for the indicated number of mice; significance determined using two-way ANOVA.

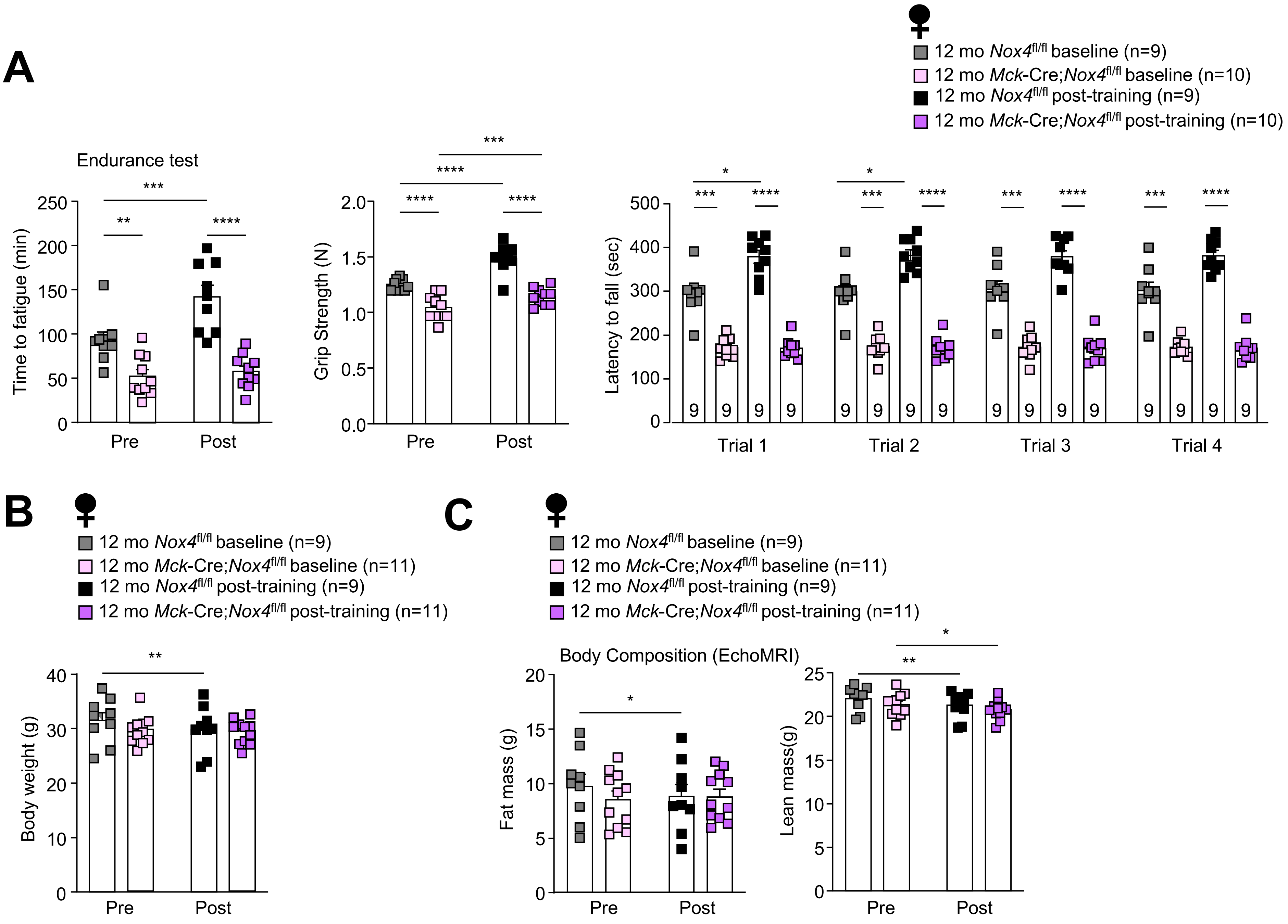


Related to Figure 6

**Figure S16**

***Figure S16 . NOX4 is required for the enhanced muscle function in responses to exercise in female mice – Related to Fig. 6e.*** *Nox4*$^{fl/fl}$ and *Mck*-Cre;*Nox4*$^{fl/fl}$ female male mice were fed a standard chow diet (4.8% fat) for 12 months and underwent basal and post exercise training (5 weeks with progressively increasing running speeds each week) measurements of **a**) endurance, grip strength and locomotor activity (rotarod). **b**) Body weights, **c**) body composition (EchoMRI) pre and post exercise training. Results shown means ± SEM for the indicated number of mice; significance determined two-way ANOVA.
